# Differing effects of size and lifestyle on bone structure in mammals

**DOI:** 10.1101/2020.08.13.249128

**Authors:** Eli Amson, Faysal Bibi

**Affiliations:** Museum für Naturkunde, Leibniz-Institut für Evolutions- und Biodiversitätsforschung, Invalidenstraße 43, 10115 Berlin, Germany

**Keywords:** Bone structure, Constructional morphology, Lifestyle convergence, Mammals, Size effect

## Abstract

The skeleton is involved in most aspects of vertebrate life history. Previous macroevolutionary analyses have shown that structural, historical, and functional factors influence the gross morphology of bone. The inner structure of bone has, however, received comparatively little attention. Here we address this gap in our understanding of vertebrate evolution by quantifying bone structure in appendicular and axial elements (humerus and mid-lumbar vertebra) across therian mammals (placentals + marsupials). Our sampling captures all transitions to aerial, fully aquatic, and subterranean lifestyles in extant mammal clades. We found that mammalian inner bone structure is highly disparate. We show that vertebral structure mostly correlates with body size, but not lifestyle, while the opposite is true for humeral structure. The latter also shows a high degree of convergence among the clades that have acquired specialised lifestyles. Our results suggest that radically different extrinsic constraints can apply to bone structure in different skeletal elements.

## Introduction

The skeleton in vertebrates is involved in many important biological roles, such as supporting body weight, locomotion, and feeding (Basu et al., 2019; Chirchir et al., 2015; Shefelbine et al., 2002). Numerous investigations of the gross morphology of skeletal elements have revealed complex relationships with ecology or life history (Bardua et al., 2020; Navalón et al., 2019). In contrast, the macroevolutionary analyses of bone structure, namely the distribution of bone tissue within a skeletal element, are scarce, creating a major impediment to our understanding of vertebrate evolution. Bone structure differs from gross morphology in its high degree of plasticity. Indeed, it was suggested as early as the late 19^th^ century that bone structure can adapt to the mechanical loads applied to skeletal elements throughout life (Kivell, 2016) (Wolff’s law, or *bone functional adaptation*). A particular phenotype is argued to be the result of structural, historical (i.e., phylogenetic), functional, and environmental factors (Briggs, 2017; Cubo et al., 2008; Seilacher, 1970). This was for instance evidenced for the gross morphology of the appendicular and axial skeleton, with comparative analyses of the forelimb (Fabre et al., 2015) in musteloids and mandible (Raia et al., 2010) in ungulates. The extent to which lifestyle, phylogenetic heritage, body size, and other structural factors influence bone structure at a broad macroevolutionary scale is poorly understood.

This study investigates bone structure and its correlates across therian mammals (placentals + marsupials). Ranging from ca. 2 grams for an Etruscan shrew and up to 170 tonnes for a blue whale, mammals occupy the depths of the ocean to aerial heights. The evolutionary history of mammals is marked by a series of diversification events, resulting in the acquisition of various specialised lifestyles as early as the Jurassic, with early mammaliaforms (Grossnickle et al., 2019). However, it is assumed that each large mammalian clade, the marsupials and placentals in particular, stems from small-sized insectivorous or omnivorous ancestors that diversified independently after the Cretaceous Terrestrial Revolution (Grossnickle et al., 2019). The subsequent evolution of each mammal clade saw transitions to highly specialised lifestyles strongly departing from those reconstructed for these Cretaceous forms. Arguably the most extreme are the aerial, fully aquatic, and subterranean lifestyles which were each acquired convergently on several independent occasions in marsupials and placentals (Ellerman, 1956; Howell, 1930; Jackson and Thorington, 2012).

We investigate the correlation between bone structure and the species’ size and lifestyle in a phylogenetically informed context. We focused our analyses on lumbar vertebrae and the humerus, these elements being generally conserved (and not vestigial) across mammals, and having been previously associated with locomotor adaptations (Buchholtz, 2001; Fabre et al., 2015; Jones and Pierce, 2016). Since mammals share broadly similar ontogenetic trajectories (compared to other tetrapods like anurans (Bardua et al., 2020)) and bony tissue types (Currey, 2003), a low disparity in bone structure across the clade should be interpreted as indicative of preponderant structural constraints. Strong phylogenetic constraints (or inertia) should in turn be associated with a strong phylogenetic signal (but see Blomberg et al. 2003). On the other hand, preeminent functional influence is expected to entail clear correlation between the traits’ distribution and lifestyle. In a similar manner, body size influence can be investigated examining the correlation of the traits and a size proxy. The diversity of mammals’ bone structure is here captured by quantifying humeral and vertebral traits for highly specialised taxa (aerial, fully aquatic, and subterranean), their terrestrial sister-groups (TSG), and more distantly related terrestrial species. Finally, we address the evolvability of bone structure. While evolvability can be generally defined as the propensity to produce evolutionary novelties, the concept can take different meaning depending on the evolutionary scale of interest (Hu and Albertson, 2016). We test for the presence of “evolutionary predispositions (Parins-Fukuchi, 2020)” by comparing these TSG to more distantly related terrestrial mammals.

## Results

### Lifestyle transitions in mammals

All reconstructed transitions to a specialised lifestyle on the tree are supported by a high posterior probability. These include no reversions (Fig. 1). From an ancestrally non-specialised lifestyle, transition events led to two aquatic convergences, cetaceans and sirenians; 13 subterranean convergences, eight in rodents, two in afrotheres, one in xenarthrans, and one in marsupials; and seven aerial convergences, three in marsupials, two in rodents, and one each in archontans, and chiropterans. The sister-groups of each of the marsupial clades that have acquired gliding are arboreal, and are hence excluded from the subsequent analyses (see Methods). The latter will consequently recognise only one acquisition of the aerial lifestyle in marsupials, and five aerial convergences in total (see also Supplementary Methods S3).

**Fig. 1.**
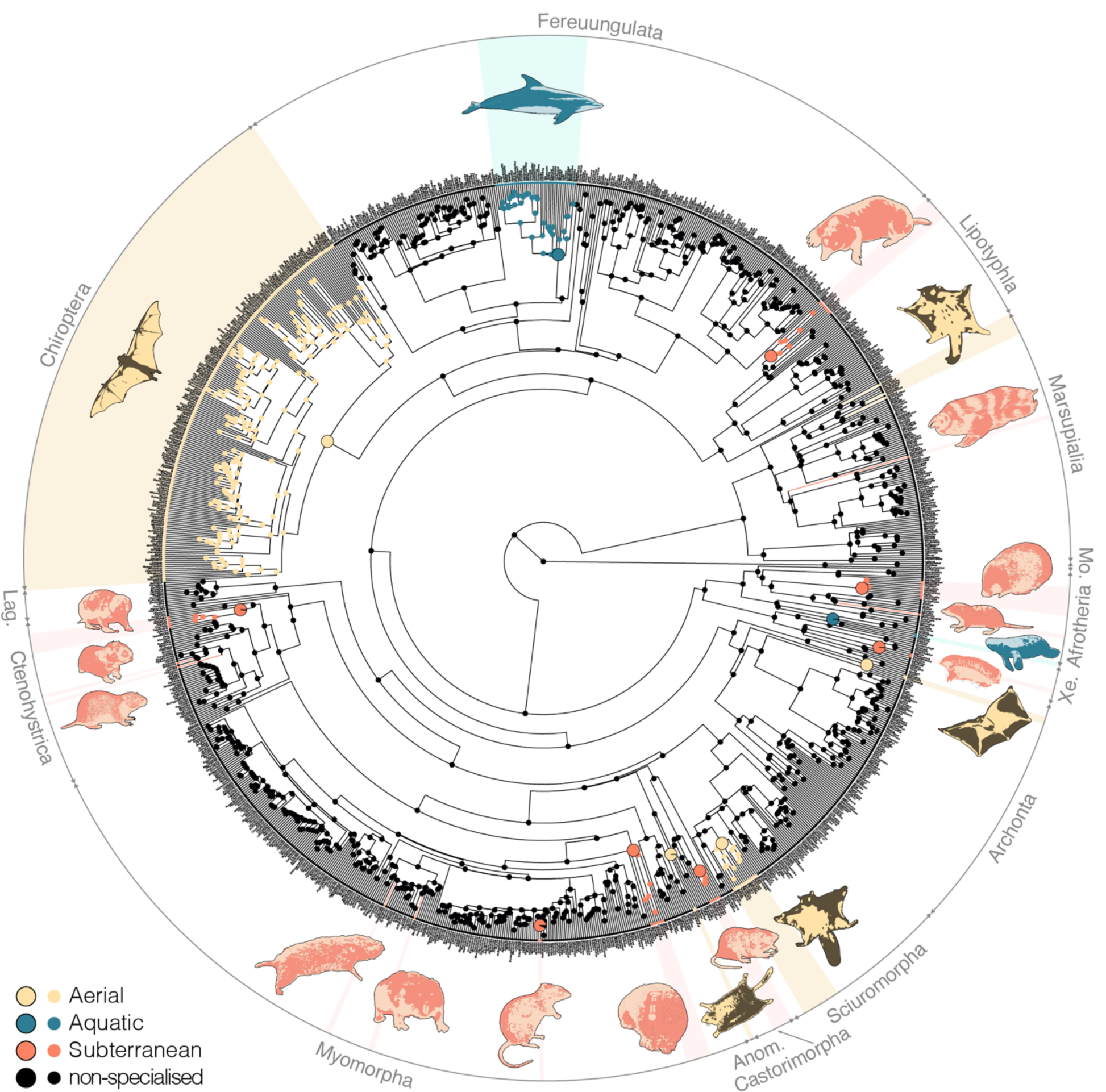
Lifestyle transitions among mammals. Colours correspond to the specialised lifestyles, i.e., aerial, aquatic, and subterranean; black corresponds to ‘non-specialised’ lifestyles. States at the nodes are reconstructed with stochastic character mapping. Lifestyle transitions were emphasised with larger nodes for clades and coloured branches for single taxa. Each silhouette represents an independent acquisition of one of the three specialised lifestyles. Abbreviations: Anom., Anomaluromorpha; Lag., Lagomorpha; Mo., Monotremata; Xe, Xenarthra.

### Size effect and lifestyle signal

Body size affects preponderantly the vertebral bone structure among mammals. Mean vertebral global compactness (Cg) and vertebral centrum trabecular architecture (Connectivity and bone fraction, BV/TV) show a strong positive correlation with size, even when lifestyle is accounted for, as indicated by phylogenetically informed ANCOVAs (pANOVAs, *p*-values < 0.0001) and phylogenetically informed regression of the parameter against a size proxy (pANOVAs, *p*-values < 0.0001, pseudo R^2^ = 0.38-0.59; Fig. 2a-c; Supplementary Results S1). Some small-sized taxa display surprisingly few (if any) trabeculae in the core of their centrum (low Connectivity; Fig. 2a, 3a), which therefore presents a low bone fraction (BV/TV; Fig. 2b). The latter conclusion can be extended to the whole vertebra, as their mean vertebral Cg is also particularly low (Fig. 2c). As they increase in size, mammals of all lifestyles show a similar increase of vertebral Cg, Connectivity, and BV/TV (Figs. 2, 3b). Accordingly, none of the vertebral traits display a lifestyle signal (Fig. 4a-c) with three exceptions (also found when accounting for size effect and when the terrestrial sample is pruned to match the size of the specialised taxa): the aquatic taxa depart from the other lifestyles in having a higher mean vertebral Cg (pANCOVAs *p*-values < 0.036) and Connectivity (*p*-values < 0.018), and the subterranean taxa differ from the terrestrial and aerial taxa in featuring a higher Connectivity (*p*-values < 0.010).

**Fig. 2.**
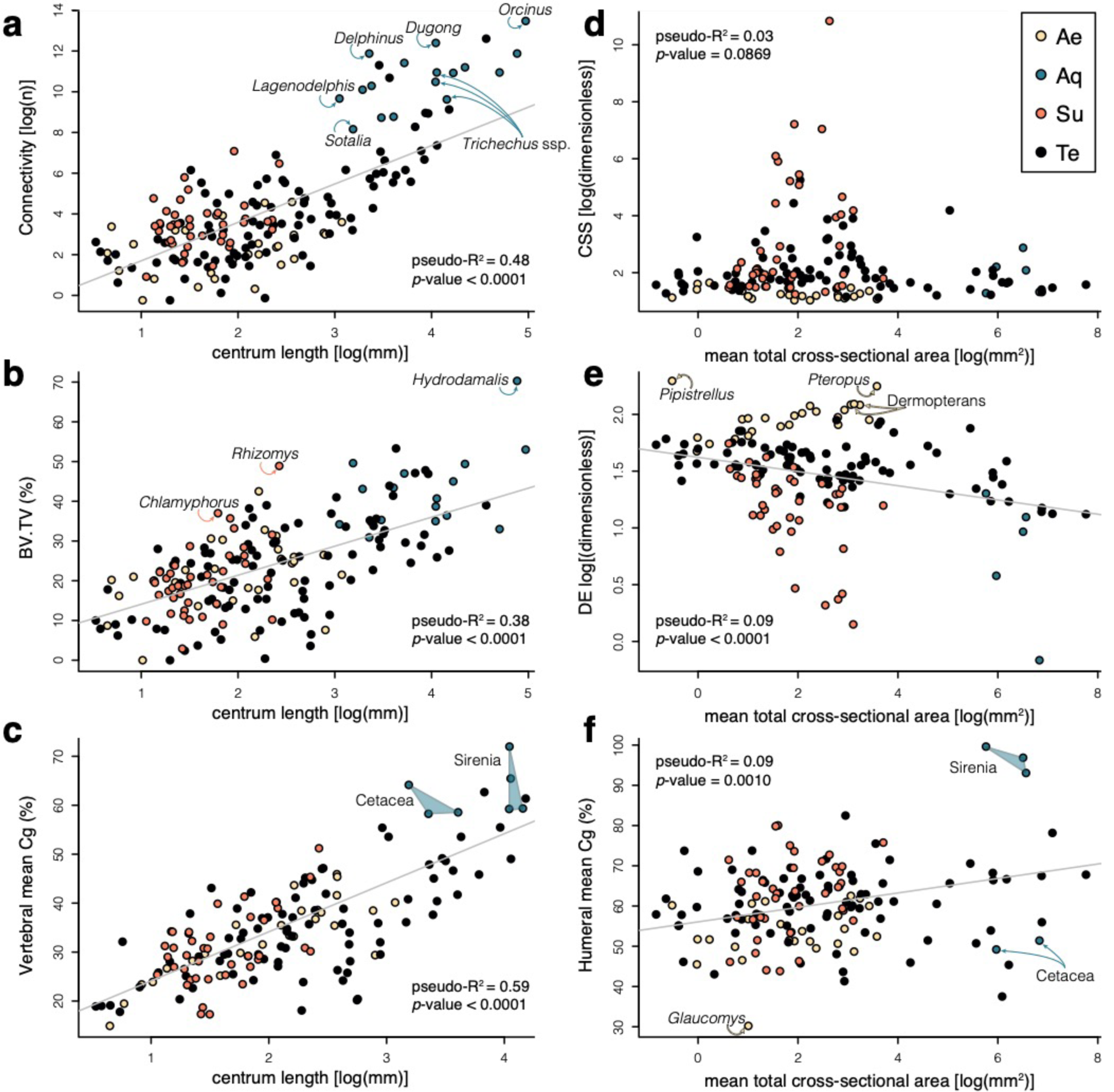
Size effect accounting for phylogeny on the studied bone structure parameters across lifestyles. Vertebral (**a**-**c**) and humeral (**d**-**f**) parameters are plotted against a body size proxy. Colours and black indicate aerial (Ae), aquatic (Aq), subterranean (Su), and terrestrial (Te) lifestyles.

**Fig. 3.**
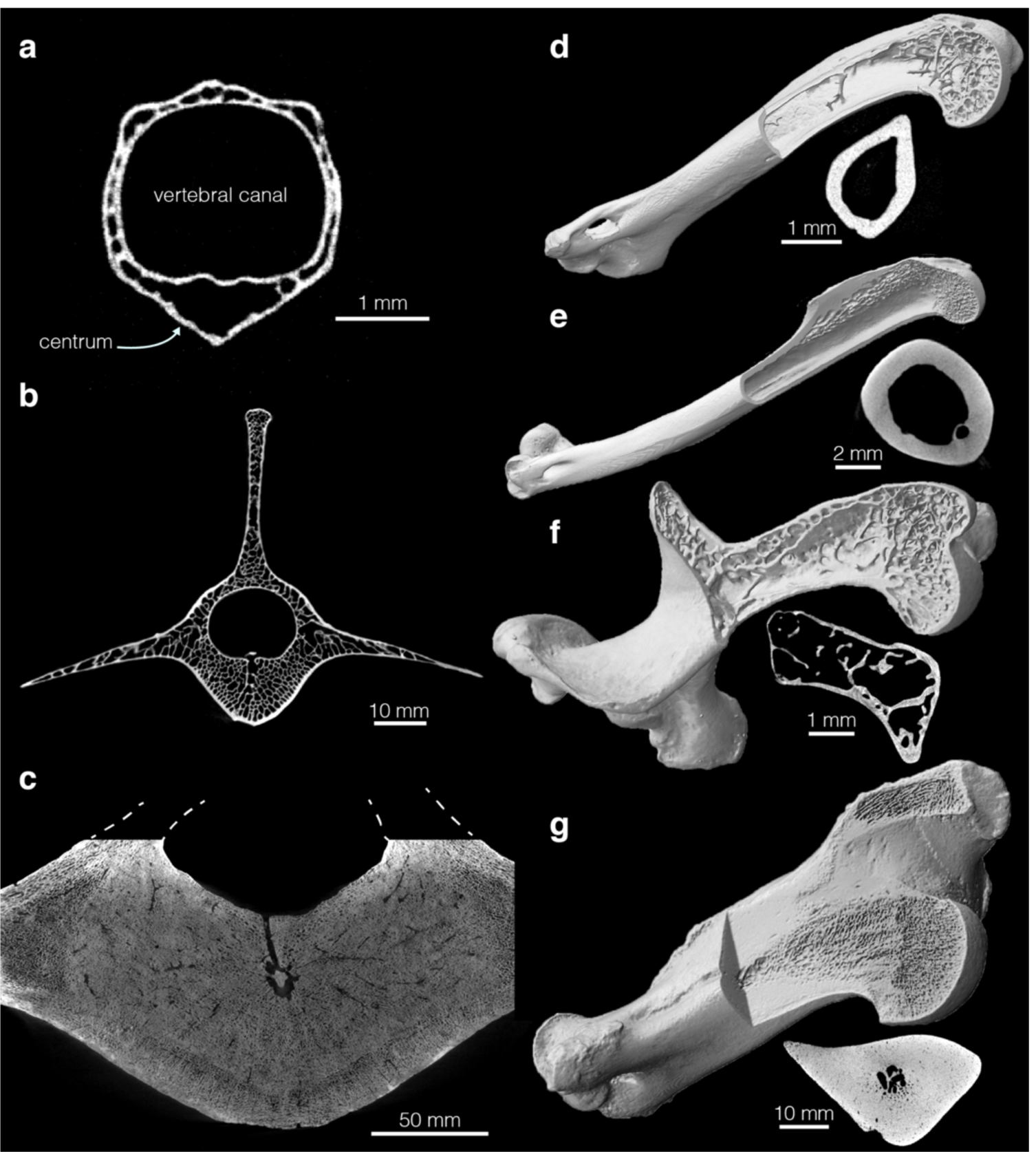
Disparity of vertebral and humeral structure among mammals. Transverse cross-section of the lumbar vertebrae at centrum’s mid-length (dorsal towards the top) for (**a**) Hose’s pygmy flying squirrel (*Petaurillus hosei*; NHMUK ZD 1900.7.29.26), (**b**) fallow deer (*Dama dama*; ZMB_Mam_94752), (**c**) Steller’s sea cow (*Hydrodamalis gigas*; MNHN_AC_1919-48). Humerus 3D rendering (not to scale) and midshaft cross-section for (**d**) the long-eared gymnure (*Hylomys megalotis*; NHMUK ZD 1999.47), (**e**) Sunda colugo (*Galeopterus variegatus*; ZMB_Mam_69096), (**f**) southern marsupial mole (*Notoryctes typhlops*; ZMB_Mam_35694), (**g**) dugong (*Dugong dugon* ZMB_Mam_69340).

**Fig. 4.**
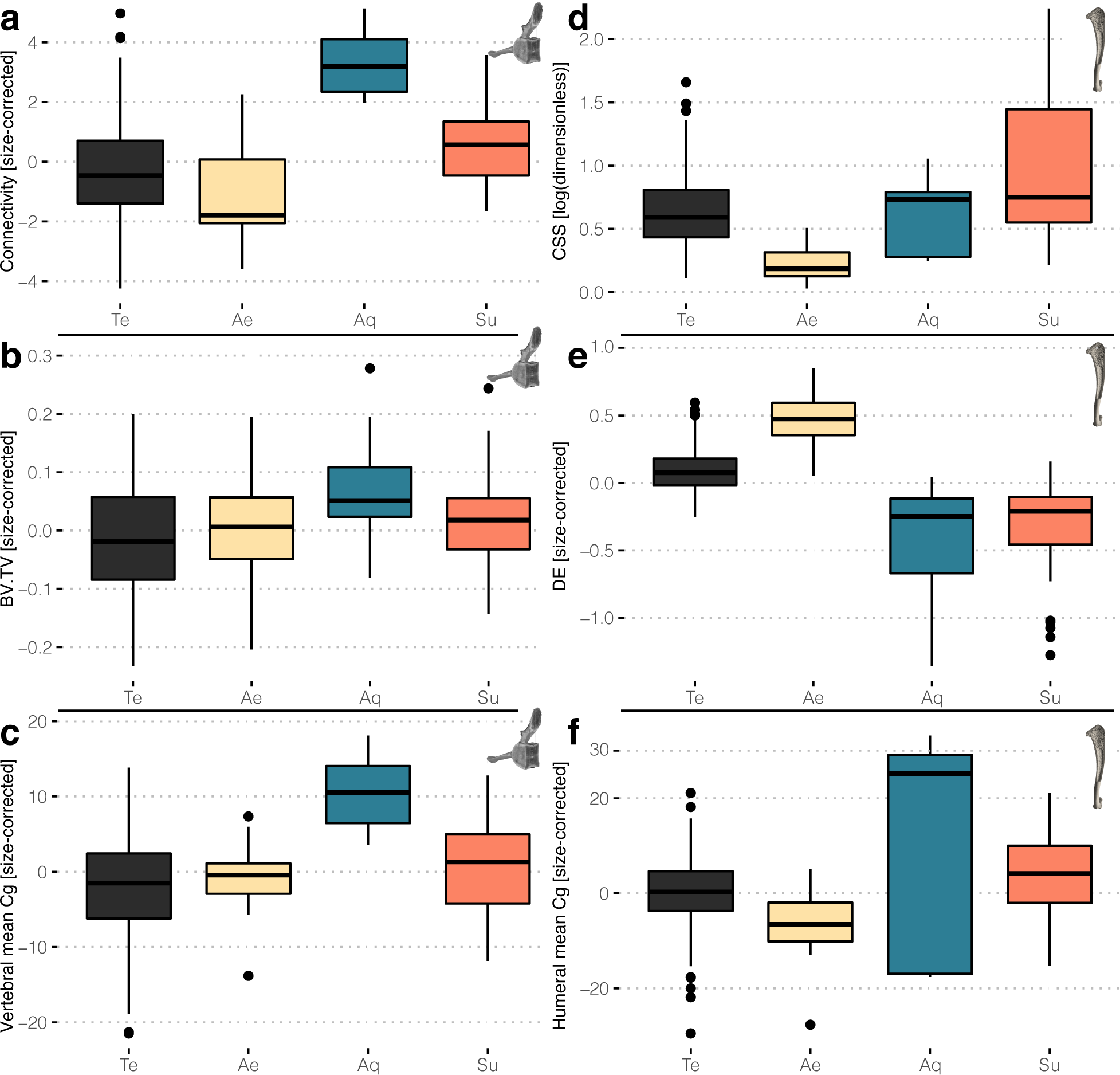
Differences among lifestyles (or the lack thereof) in the studied bone structure parameters. Boxplots (centre line, median; box limits, upper and lower quartiles; whiskers, 1.5 times interquartile range; points, outliers) depicting the distribution of (**a**-**c**) vertebral parameters and (**d**-**f**) humeral parameters. Abbreviations: Ae, aerial; Aq, aquatic; Te, terrestrial; Su, subterranean.

Conversely, there is no preponderant size effect on the investigated humeral parameters, namely the mean humeral Cg, cross-sectional shape (CSS), and diaphysis elongation (DE) (pANOVAs parameter ∼ size, all pseudo R^2^ < 0.10; Fig. 2d-f; Supplementary Results S1). Only DE seems to consistently scale (with weak negative allometry) across lifestyles (pANCOVA, *p*-value < 0.001; Fig. 2e). The correlation between mean humeral Cg and the size proxy (pANCOVA, *p*-value = 0.018) is mostly driven by the high values of the large-sized sirenians (Fig. 2f). A clear lifestyle signal is found in the humeral structure, including when the size effect is accounted for and when the terrestrial sample is pruned to match the size of the specialised taxa (Fig. 2d-f; Fig. 4d-f; Supplementary Results S1). Aerial taxa (Fig. 3e) differ from subterranean (Fig. 3f) and terrestrial taxa (Fig. 3d) in featuring a diaphysis that is more elongate (higher DE; pANCOVA *p*-values < 0.001) and also more circular in cross-section (lower CSS; *p*-values < 0.002). Furthermore, subterranean taxa differ from terrestrial taxa in featuring a lower DE (*p*-value < 0.001) and higher CSS (*p*-value < 0.024). Finally, aquatic taxa (Fig. 3g) differ from aerial taxa in their more compact humeral structure (higher mean humeral Cg, *p*-value = 0.047), but one should bear in mind that the two lifestyles also differ in their body sizes. Moreover, a great disparity is observed in the mean humeral Cg values of aquatic taxa, due to the outstandingly high values of sirenians (Figs. 2f, 3g). Aerial and subterranean taxa were also found as having lower DE values than the aquatic ones (*p*-value < 0.01), but the lack of size overlap between the groups prevents a strict exclusion of the size effect.

### Convergence analysis

Examination of the strength and direction of the convergence in the studied parameters’ evolution allows to better understand the modalities by which each specialised lifestyle was acquired among mammals. For the subterranean lifestyle, acquired by 13 clades, the Connectivity (corrected for size effect) do show overall convergence (C1 = 0.52, *p*-value < 0.01) with twelve clades increasing their mean value compared to the reconstructed value of their respective ancestral nodes (five exceeding the corresponding 95% confidence intervals [95CIs]; Fig. 5a). DE on the other hand does not show strong overall convergence (C1 = 0. 24, *p*-value = 0.82; using size-corrected values), but twelve clades decreased their values (six exceeding the 95CIs; Fig. 5b). No strong overall convergence is found in CSS either (C1 = 0.25, *p*-value = 0.78), and seven clades show an increase of their values (four exceeding the 95CI; Fig. 5c). The distribution of all traits, including the other, non-converging ones, can be found in Supplementary Results S2.

Among aerial taxa, overall convergence is the clearest for CSS (C1 = 0.71, *p*-value < 0.001), with all five clades decreasing their values compared to their respective ancestral nodes (three exceeding the 95CIs). DE also shows overall convergence (C1 = 0.41, *p*-value = 0.040), with all clades showing an increase of their values (four exceeding the 95CIs). Humeral mean Cg does also converge among aerial clades (C1 = 0.72, *p*-value = 0.002), but with only three clades showing a decreasing (two exceeding the 95CIs). While Connectivity does not show clear evidence of convergent evolution (C1 = 0.47; *p*-value = 0.0495), four out of five clades feature a decrease of their values (one exceeding the 95CI).

**Fig. 5.**
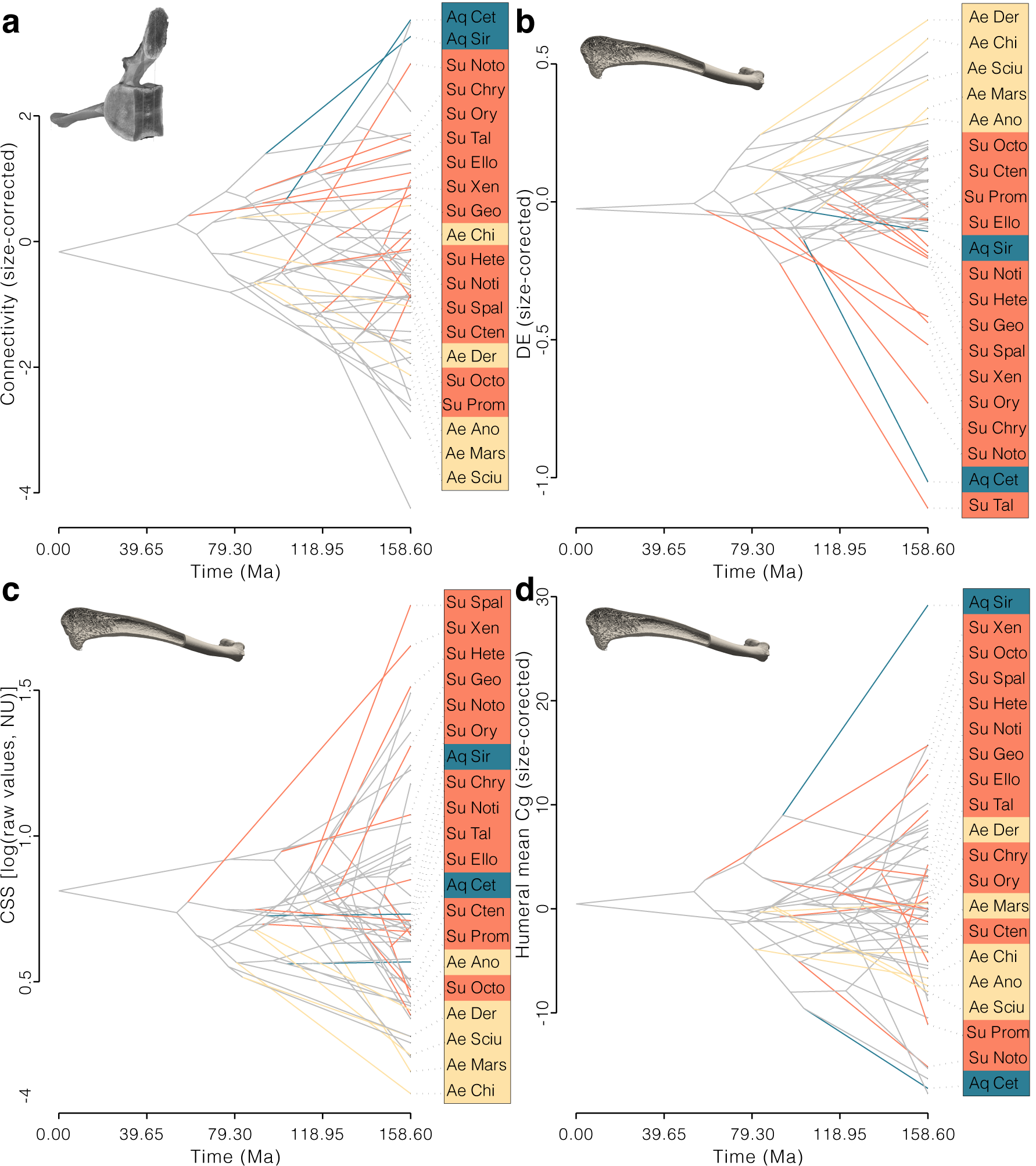
Trait convergence among mammalian lifestyles. Phenograms depicting reconstructed evolution of the vertebral centrum Connectivity (**a**) and humeral diaphysis elongation (**b**), midshaft cross-sectional shape (**c**), and mean global compactness (**d**). Note the number of branches leading to the specialized clades (coloured) evolving in the same direction for converging traits (e.g., in (**a**) twelve out of 13 subterranean clades increased their Connectivity when compared to their direct ancestor). Clade abbreviations: see Supplementary Table S1.

For aquatic clades, only BV/TV (size-corrected) shows a clear overall convergence (C1 = 0.97; *p*-value = 0.010), with both sirenians and cetaceans increasing when compared to their respective ancestral nodes (neither exceeding the 95 CIs). One parameter is actually diverging between the two aquatic clades: the humeral mean Cg increased in sirenians while it decreased in cetaceans (both exceeding the respective 95CIs; Fig. 5d). The absence of overall convergences for the vertebral mean Cg might be due in part to the size correction we used for this parameter, which weakens the signal (uncorrected values fall beyond most of other lifestyle’s values). For this parameter both clades show an increase of their values (only cetaceans exceed the corresponding 95CI). The same is true for the Connectivity (C1 = 0.88; *p*-value = 0.052), for which both clades feature an increase of their values (both exceeding the 95CIs).

### Evolvability analysis

All converging traits differ between the specialised and terrestrial lifestyles (Fig. 6). But specialised clades’ respective terrestrial sister-groups (TSG) do not differ from the other (more distantly related) terrestrial taxa we sampled for any of the investigated traits. It is only for the Connectivity (vertebral centrum) in subterranean clades and CSS in aerial clades that the respective TSG tend to fall between the more distantly terrestrial and specialised clades (Fig. 6a, f). The opposite is true for CSS in subterranean clades.

**Fig. 6.**
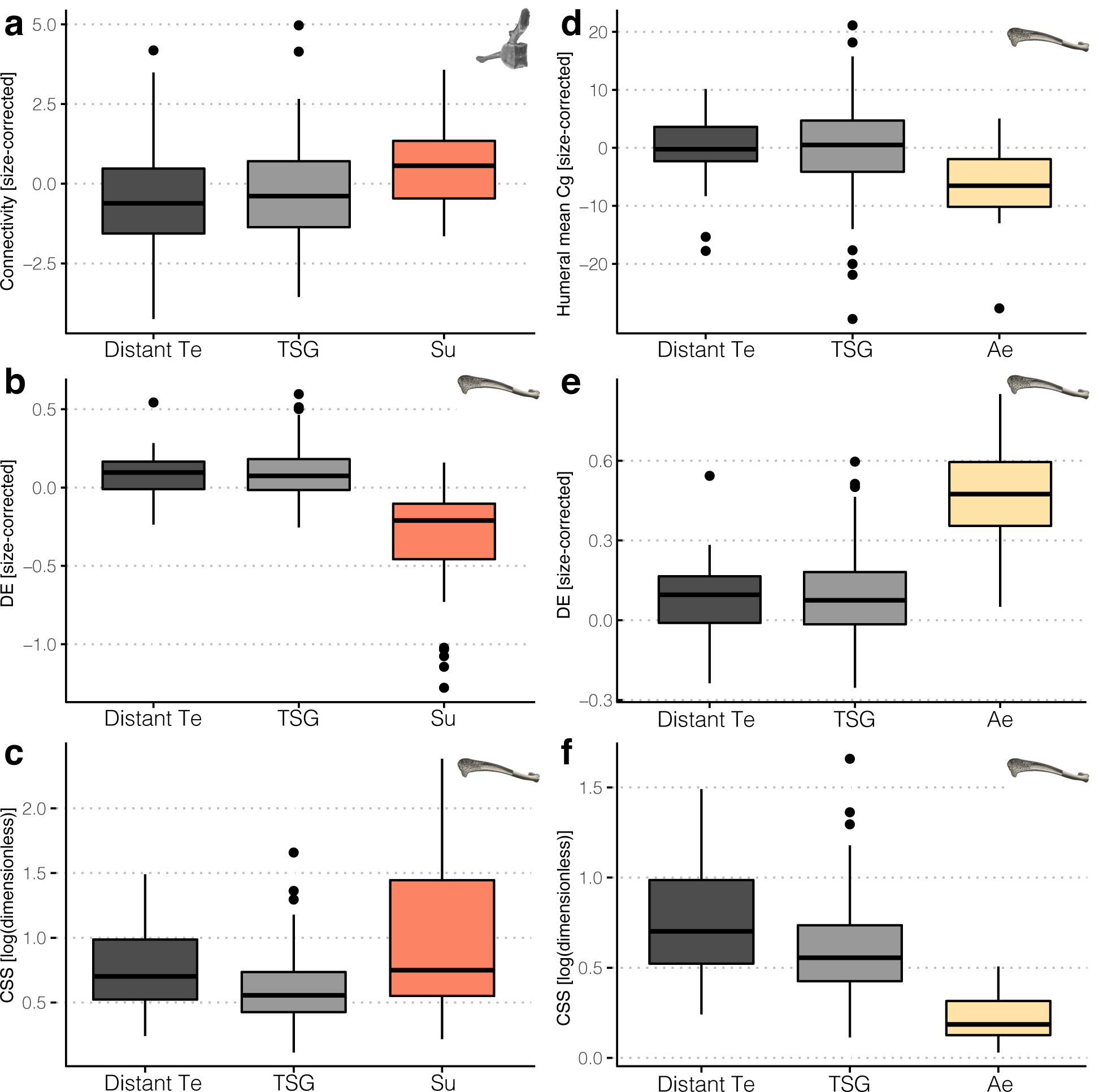
Evolvability analysis. The traits found as converging were compared among subterranean (**a**-**c; ‘**Su’) and aerial (**d**-**f**, ‘Ae’) clades, their respective terrestrial sister-groups (‘TSG’), and more distantly terrestrial taxa (‘Distant Te’). Boxplots definition: centre line, median; box limits, upper and lower quartiles; whiskers, 1.5 times interquartile range; points, outliers.

### Phylogenetic signal

A significant phylogenetic signal was found for all traits in terrestrial taxa when all species were analysed independently (lambdas = 0.25-0.88; *p*-values < 0.022; Supplementary Results S3). However, when the members of the TSG are aggregated to sample the mammalian tree more uniformly, none of the traits were showing a significant phylogenetic signal except for the Connectivity and DE (lambdas = 0.84-0.85; *p*-values < 0.03).

## Discussion

All bone structure parameters herein investigated are correlated to body size and/or lifestyle. Controlling for the potential effects of lifestyle and phylogeny, size is clearly correlated with all vertebral traits. with the notable exception of the aquatic lifestyle, analyses of covariance and quantification of the convergence suggest that lifestyle exerts little influence on these traits. As for several gross morphological traits of the vertebral column in (semi-)aquatic mammals (Buchholtz, 2001; Jones and Pierce, 2016), we found that vertebral inner structure of aquatic species differ from that of their terrestrial relatives (beyond what can be expected from a size effect alone; see below). However, the other specialised lifestyles we have investigated cannot be clearly associated with a particular vertebral phenotype. The opposite is true for the traits capturing the humeral structure we have investigated: little or no influence of size was detected, while each specialised lifestyle is associated with a specific phenotype departing from that observed in closely related terrestrial taxa.

Our data emphasises an unsuspected diversity in vertebral inner anatomy of therian mammals, with, for example, an overall bone fraction ranging from ∼15 to 72% (Fig. 3; Supplementary Table S2). High disparity in humeral structure was also observed, with the bone fraction ranging from ca. 30 to over 99% for the humeral diaphysis. Those are among the most extreme values ever quantified for amniotes (Canoville and Laurin, 2010; Houssaye et al., 2013), which lends support to the conclusion that constraints (such as ontogenetic ones) are not acting particularly strongly on the bone structure of mammals. Furthermore, we found only weak phylogenetic signal in the investigated traits when mammals are sampled uniformly. While that does not strictly entail the absence of phylogenetic constraints (Blomberg et al., 2003), it indicates that phylogeny explains very little of the observed trait distributions. Taken together, these considerations suggest that size (in vertebrae) and lifestyle (in humerus) should be viewed as the main factors impacting bone structure in the mammalian postcranium.

The influence of body size on the vertebral structure is most compelling when examining the lower extreme of the mammalian spectrum: the mid-lumbar vertebra of small mammals is often delicate: the overall compactness (mean vertebral Cg) than can be as low as 15% and no or few trabeculae are observed the middle of the centrum (Figs. 2, 3). This has to be contrasted with the structure commonly observed in larger terrestrial mammals: a centrum entirely filled with spongy bone, and an overall compactness of ca. 30-50%. A more robust construction of the vertebrae with increasing size is therefore the rule in mammals, as all vertebral traits investigated scale with positive allometry (Fig. 2). A similar conclusion was reached for the dorsal vertebrae of squamates (Houssaye et al., 2010). Several vertebral gross morphological traits were shown to scale with allometry in terrestrial mammals (Halpert et al., 1987; Jones, 2015; Jones and Pierce, 2016). These have been importantly correlated to an increased stiffness of the vertebral column with larger sizes. Both gross morphology and structure are hence in accordance with an area:volume scaling (Biewener, 2005), which predicts that stresses will increase with body mass because the strength of the vertebral column only scales with cross-sectional area. That biomechanical interpretation should be tempered regarding the mean vertebral Cg, however, because the vertebral canal seems to scale with negative allometry (see, e.g., Fig 3a). While this was not assessed by our analysis, this seems to be confirmed by the fact that the spinal cord weight scales with negative allometry in mammals (MacLarnon, 1996). However, this explanation cannot be brought forward for the negative allometry we recovered for the traits related to the vertebral centrum, which were acquired in a volume of interest placed in its centre. In mammals, the centrum is expected to mainly withstand axial compression during bending of the vertebral column (Slijper, 1946). Using a finite element analysis of the human centrum loaded in compression, it was shown that relative loads shared by the cortex and trabecular bone depend on the distance from the cranial and caudal ends of the centrum (Eswaran et al., 2005): the greatest fraction of load taken by the cortex (up to 54% of the total load) was found in the middle of centrum. Our data suggest that under a certain body size, loads could be low enough to be mostly (if not entirely) withstood by non-trabecular tissues.

The aquatic lifestyle is the class for which we found the clearest association with specific vertebral structures (also valid for the humeral structure, see below). This could be ascribed to the drastically different medium in which cetaceans and sirenians locomote, relieving the vertebral column from the loads associated with a quadrupedal stance and terrestrial locomotion (notably due to Archimedes’ principle). Indeed, other constraints are associated with aquatic locomotion, and two major bone structure adaptations are assumed to relate to buoyancy and trim control during swimming (Ricqlès and Buffrénil, 2001). Bone mass increase (BMI; classified as non-pathological pachyostosis and/or osteosclerosis) has been reported in sirenians, and interpreted in the light of their shallow diving habits (Buffrénil et al., 2010). Conversely, qualitative assessments of cetaceans’ vertebral structure have reported bone mass decrease, notably due to thinner cortices (Houssaye et al., 2016). However, previous quantification of the 2D bone structure of the centrum did not reveal considerable differences of their bone fraction with their terrestrial relatives (Dumont et al., 2013; Houssaye et al., 2014). This is also what is indicated by the ANCOVA we performed on BV/TV (Fig. 2b; Supplementary Results S1). This trait was nonetheless found as converging in our analysis, with both clades having acquired greater values when compared to the reconstructed ancestral ones (Supplementary Results S2). Our analysis more clearly revealed that the overall vertebral bone fraction (mean vertebral Cg) is high in both sirenians and cetaceans (Fig. 2c). Because both parameters are affected by positive allometry, and because sirenians and cetaceans are among the largest mammals included in the dataset, it is difficult to safely associate high mean vertebral Cg with the aquatic lifestyle (as taking into account size as covariate or using size-correction will be least accurate for extreme values). Based on our analysis, there is nevertheless no reason to acknowledge overall vertebral BMI in sirenians and not in cetaceans. The convergence analysis even showed that the latter more sharply increase from the corresponding reconstructed ancestral value. Both sirenians and cetacean might hence share similar functional constraints upon overall vertebral bone fraction. The conclusion has to be tempered by the extreme case of the Steller’s sea cow (*Hydrodamalis gigas*; Fig. 3c), its centrum’s bone fraction exceeding 70%. Such a value, unparalleled in extant mammals, confirms that with the recent extinction of this species we lost a truly exceptional component of mammalian diversity (Domning, 1976). We also found that the number of trabeculae (the other investigated vertebral trait, Connectivity) is greater in sirenian and cetaceans (Figs. 2a, 4a, 5a). This agrees with Dumont et al. (2013)’s similar observation regarding cetaceans. They interpreted this feature as potentially related to protecting the vertebra from accumulating fatigue microfractures, assumed to occur during their locomotion, which entails high frequency cycles of tension and compression exerted on the centra. Our results concur quite well with this interpretation, with manatees and coastal cetaceans featuring relatively less trabeculae in their centrum (accounting for the size effect on Connectivity) than the more active pelagic cetaceans and dugong (Kojeszewski and Fish, 2007) (Fig. 2a).

We found only one other vertebral parameter that showed a correlation with the investigated lifestyles: Connectivity was found as greater in subterranean species (Figs. 2a, 4a, 5a; Supplementary Results S1). This can be interpreted as indicative of greater axial compressive loads on the vertebral column (Smith and Angielczyk, 2020). Such greater loads could be expected to be associated to subterranean lifestyle: during the propulsion phase of digging, the vertebral column transmits the soil reaction force (though the hind limbs that anchor body to the substrate (Gasc et al., 1986)). The strength and stiffness of trabecular bone being mostly due to its bone fraction (Nazarian et al., 2008), BV/TV (and to a lesser extent the mean vertebral Cg as well) should similarly be expected to be greater in subterranean species, which was not clearly featured by our data. Nevertheless, some subterranean taxa we have sampled, such as the pink fairy armadillo (*Chlamyphorus truncatus*), were marked by outstandingly high BV/TV values (Supplementary Results S2). For the other subterranean species, BV.TV that is not particularly high but with higher Connectivity should entail thinner trabeculae (trabecular thickness was not assessed here). The number of trabeculae rather than their thickness is responsible for the strength of trabecular bone (Guo and Kim, 2002). Our results hence suggest that some subterranean clades might have developed a vertebral column structure capable of withstanding relatively greater loads without increasing its overall bone content, except for some cases for which additional strengthening might be required.

Sharply differing from the vertebral bone structure, humeral structure correlates with each investigated lifestyle rather than body size. The cross-sectional shape (CSS) differentiates well the aerial and subterranean clades (Figs. 2d, 4a, 5c). Each of the five aerial clades convergently acquired a more circular diaphysis in cross-section (lower CSS values; Fig. 3e). Bats have the most circular diaphysis. The opposite trend is associated with the subterranean lifestyle, with in particular the spalacids (blind mole-rats and allies), heterocephalids (naked mole-rat and allies), geomyidae (pocket gophers), and the pink fairy armadillo that acquired a highly elliptical cross-section of the humeral diaphysis (Fig. 3f). Our assessment of CSS therefore illustrates quite well the spectrum of constraints acting on the humeral diaphysis, ranging from round cross-sections (with relatively thin cortex, see below) that maximise resistance to torsional stresses in volant taxa (Swartz et al., 1992) to the more elliptical cross-sections that better withstand uniaxial bending loads associated with digging (Amson and Nyakatura, 2018). The low CSS in the non-volant aerial mammals, i.e., the four gliding clades, could be reflective of the need to resist torsional loads as in bats and/or to a multidirectional bending environment, which is assumed in the case of the similarly rounder cross-sections of non-aerial, arboreal mammals (Patel et al., 2013). The low level of convergence we found for CSS among subterranean clades likely emphasises the disparity of fossorial behaviours, which for instance include scratch- and humeral rotation digging (Kley and Kearney, 2007). The talpids, who exemplify the latter digging style, were not characterised by a high CSS (and did not acquire higher values relative to the ancestral value; Fig. 5c). However, their humeral structure is clearly specialised, which is for instance demonstrated by their more elliptical medullary cavity (Meier et al., 2013). The humerus of other mammalian fossors were shown to feature relatively high values of cross-sectional parameters such as the second moment of area or the polar section modulus (Hedrick et al., 2020; Kilbourne and Hutchinson, 2019).

The humeral diaphysis elongation (DE) also differentiates well the aerial and subterranean clades (Figs. 2e, 4a, 5b), with elongate bones for the former (Fig. 3e) and stouter bones for the latter (Fig. 3f). Aquatic taxa also have a stout humerus (falling in the range of subterranean clades), but that is mostly expressed in cetaceans, and convergence is hence weak in their case. As for CSS, the aerial clades are featuring the strongest convergence, with each clade increasing its value when compared to that of its respective ancestral node. While this trait scales with negative allometry for all other taxa (being the only humeral trait that shows a rather clear correlation with size), aerial species are clearly differing in showing positive allometry (Fig. 2e). This follows the expectation that wing surface should increase with positive allometry to be able to support increasing mass (Runestad and Ruff, 1995). We do however find that the smallest sampled bat (*Pipistrellus pipistrellus*) outstands drastically from this scaling in featuring an exceptionally elongate humerus. Colugos (dermopterans) are recognised as differing from other gliding mammals notably regarding the construction of their patagium, somehow approaching the chiropteran condition (Beard, 1993). It is hence not surprising that both clades feature the most elongate diaphysis of our dataset (high DE values; Fig. 5b). The average DE value is actually lower in bats than in colugos. This likely emphasises that the handwing –trivially present in gliding mammals– is a key chiropteran autapomorphy (Amador et al., 2019). Only a weak convergence was found for DE among subterranean clades (C1 = 0. 24), but it is noteworthy all clades but one (*Spalacopus cyanus*, the coruro) decreased their values when compared to their respective ancestral nodes (Fig. 5b). We interpret that pattern as indicative of the disparity of fossorial adaptations combined to the nevertheless commonality of a relatively high bending strength for the humerus of subterranean taxa. It is rather clear that a strong negative allometry affects this trait in subterranean taxa, differing from the positive or weak negative scaling relationships found for the other lifestyles (see above; Fig. 2e).

The mean humeral global compactness (Cg) in turn most clearly discriminates those bone structure adaptations that relate to buoyancy and trim control (Ricqlès and Buffrénil, 2001). The dichotomy between the sirenian osteosclerosis (Fig. 3g) and cetacean osteoporosis is compelling, and logically recovered as strongly diverging between the two clades in our analyses (Figs. 2f, 5d). The examination of this trait also allows to tackled the question of lightweightness among mammals (Dumont, 2010). The ANCOVA we performed suggests that the specialised lifestyles are not associated with differences in humeral Cg. However, the convergence analysis interestingly showed that this trait does converge among aerial clades, especially for bats, anomalurids (scaly-tailed squirrels), and gliding sciurids (squirrels) that have acquired low Cg values (Fig. 5d). There is no major difference in the bone tissue density in mammals, which has been demonstrated in particular for the humerus of bat and various rodents (Dumont, 2010). One can therefore truly recognise a tendency for bats, scaly-tailed squirrels, and gliding squirrels to have acquired a lightweight humerus. Because the humeral diaphysis of all the aerial species we sampled was basically tubular and of consistent Cg along the diaphysis, one can assimilate the mean humeral Cg to the relative cortical area (or thickness, also equivalent to the parameter K (Currey and Alexander, 1985)) The average value for these species is 53%, which falls in the upper range of the relative cortical area of volant birds (Habib and Ruff, 2008; Voeten et al., 2018).

Correlation with body size is nevertheless quite clearly present for one of the investigated humeral traits, DE. This trait naturally relates to humeral length, which has been shown to scale with weak negative allometry in mammals and tetrapods in general (Campione and Evans, 2012) (body mass ∼ humeral length^2.9^; scaling coefficient with our dataset pruned to terrestrial species: 2.8). This is consistent with the weak, negative allometry we found for DE. Our data however suggest that this allometric effect (DE ∼ body size proxy^−0.063^; dataset pruned to terrestrial species: −0.058) is minor when compared to the differences that can be imputed to lifestyle, as aerial and subterranean taxa feature conspicuously elongate or stouter bones, respectively (absolute value of coefficients > 0.26; Supplementary Results S1).

We found no ‘evolutionary predispositions’ for any of the converging traits (Fig. 6). This could entail that this test is overly simplistic. Furthermore, the taxa most closely related to the specialised species were often excluded from our sampling because of their incipient specializations (e.g., arboreal petauroids). Nevertheless, it is also possible that evolutionary predispositions are not relevant to highly plastic traits such as bone structure (Barak et al., 2011; Currey, 2003; Lanyon and Rubin, 1985; Mittra et al., 2005), which in turn can be canalised to evolve functional specializations (Lister, 2014).

In summary, the bone structure of two representative skeletal elements of the appendicular and axial skeleton, the mid-lumbar vertebra and the humerus, was found as astonishingly disparate in therian mammals. The distribution of vertebral traits was predominantly explained by mammal’s body size but not their lifestyle, while the opposite was true for humeral traits. A simple comparison between specialised species’ unspecialised sister-groups and more distantly related taxa revealed no ‘evolutionary predispositions.’ We found that 13, five (plus two counting convergences among gliding marsupials), and two clades represented by extant species respectively acquired subterranean, aerial, and fully aquatic lifestyles convergently. Those convergences are detectable in the humeral bone structure of mammals. These conclusions should be corroborated with a more exhaustive assessment of therian evolution, which could be undertaken by sampling extinct representatives of these highly specialised lifestyles, such as the gliding eomyids (Storch et al., 1996) or glirids (Mein and Romaggi, 1991) for instance.

## Materials and Methods

### Lifestyles, species, and tree

We defined specialised lifestyles as follows: fully aquatic, species that live exclusively in water (Howell, 1930); aerial, species that are able to fly or glide (Jackson and Thorington, 2012); and subterranean, species spending the greater part of their lives underground (Ellerman, 1956). We regarded the remainder of the mammalian lifestyles, including semi-aquatic or arboreal taxa, as ‘non-specialised.’ These four lifestyle classes were coded for all extant mammal genera (Fig. 1; no lifestyle variation was recognised within genera). The main timetree used for phylogenetically informed analyses was extracted from TimeTree.org (Kumar et al., 2017).

Furthermore, we acquired bone structure data for all specialised clades, as well as for the most closely related taxa of each clade that can be regarded as terrestrial, i.e., not described as (semi-) aquatic, arboreal/scansorial, or fossorial. Taxa presenting these less specialised, non-terrestrial lifestyles were excluded from the sampling because of the potential effects these lifestyles can entail on bone structure. Primary data regarding non-specialised species’ lifestyles were taken from Nowack (1999). Finally, representatives of the other terrestrial families (defined with the same criteria) were also sampled, in an endeavour to cover the diversity of extant mammals. As a result, representatives of each specialised clades, their terrestrial sister-groups (TSG), and of all other terrestrial families but two (Dinomyidae and Antilocapridae) were sampled (Supplementary Fig. S1). It was endeavoured to sample at least 5 specimens per specialised clade and its TSG. In some clades, represented by one or two species, that motivated the acquisition of repeats for the same species (e.g., the marsupial mole, *Notoryctes typhlops*). In addition to these extant species, we have extended the sampling of the sirenians (represented by two extant genera) with the recently extinct Steller’s sea cow (*Hydrodamalis gigas*), given that skeletal remains with a preservation comparable to that of extant species were available. This amounts to 190 museum specimens, representing 182 species (Supplementary Table S1). The main timetree was pruned to the subset of sampled species and missing species were added by hand (see Supplementary Methods S1). Lifestyle classification of all genera and timetree with 182 sampled species can be found on figshare (doi: 10.6084/m9.figshare.12600440).

### Phenotypic data

Several traits were acquired to capture the bone structure properties of the whole humerus and whole mid-lumbar vertebra (in consistency with Dumont et al. 2013). Micro-Computed Tomography (µCT) data were acquired for both skeletal elements. Scanning was performed with a Phoenix nanotom (General Electric GmbH Wunstorf, Germany), a FF35-CT-System (YXLON GmbH Hamburg, Germany), and HMX ST 225 (Nikon Metrology UK Ltd.). Scanning resolution was chosen to comply with the minimum relative resolution recommended for trabecular architecture analysis (Sode et al., 2008). Scans acquired by Dumont et al. (2013) were also used to expand the vertebral centrum dataset (see below). Prior to parameter acquisition, both the lumbar vertebra and the humerus were given a standard orientation using the software VG Studio Max 3.0-3.3 (Volume Graphics, Heidelberg, Germany): the bones’ mediolateral and anteroposterior/proximodistal axes were aligned along the axes of the stack. All subsequent data acquisition was performed with the software Fiji/ImageJ (Schindelin et al., 2012) (2.0). Many studies have quantified bone structure traits for long bones, and we will focus on three parameters for the humerus: CSS, DE, and mean Cg, which can be argued to reflect torsional and bending loads for the former two, as well as compressive loads (terrestrial locomotion) and buoyancy (aquatic locomotion) for the latter (Houssaye and Botton-Divet, 2018; Kilbourne and Hutchinson, 2019; Stanchak et al., 2019; Swartz et al., 1992). CSS –the ratio of maximum and minimum second moment of area (Ruff and Hayes, 1983)– was acquired at midshaft (plug-in *Slice Geometry*, BoneJ v. 1.4.; Doube et al., 2010). DE was defined as the ratio between the functional length of the humerus and the square root of the total cross-sectional area at midshaft (making this parameter dimensionless). Humeral mean Cg was acquired using the approach described in Amson (2019) (Fig. 7). In brief, slice-by-slice profiles are computed for each parameter of interest along an anatomical axis. In this case, proximodistal profiles were computed for Cg in a region spanning the middle 40% of the humerus’ functional length (to ascertain that only the diaphysis would be sampled across all studied taxa and their disparate bone morphology; Fig. 7a). Mean Cg is then computed based on the mean value of all included slices. The total cross-section area was also measured in this region for its mean value to be used as a body size proxy (see below). Fewer studies have quantified bone structure traits for vertebrae. For biomechanical reasons, the centrum and its trabeculae were previously analysed in comparative studies (Dumont et al., 2013). But it became clear that such an approach would not suffice for our dataset, as many small mammals feature a simple centrum architecture with few trabeculae. We hence also measured mean Cg in the central region of the whole vertebrae (i.e., using slice-by-slice profiles comprising all those cross-sections for which the vertebral canal is complete; Fig. 7b) to have an overall assessment of the vertebral robusticity (see similar approach based on single cross-sections by Houssaye et al., 2019). BoneJ was also used to quantify 3D trabecular architecture for cubic VOI defined to be as large as possible while being centred in the middle of the lumbar vertebra’s centrum (Smith and Angielczyk, 2020). Because few (if any) trabeculae were included in the VOI of some taxa (see Results), we refrained from further analysing trabecular parameters that would be spurious, and focused on the VOI’s BV/TV (plug-in *Volume Fraction*) and Connectivity (plug-in *Connectivity*; approximates the number of trabeculae). The anteroposterior length of the vertebral centrum was also measured to be used as a body size proxy.

**Fig. 7.**
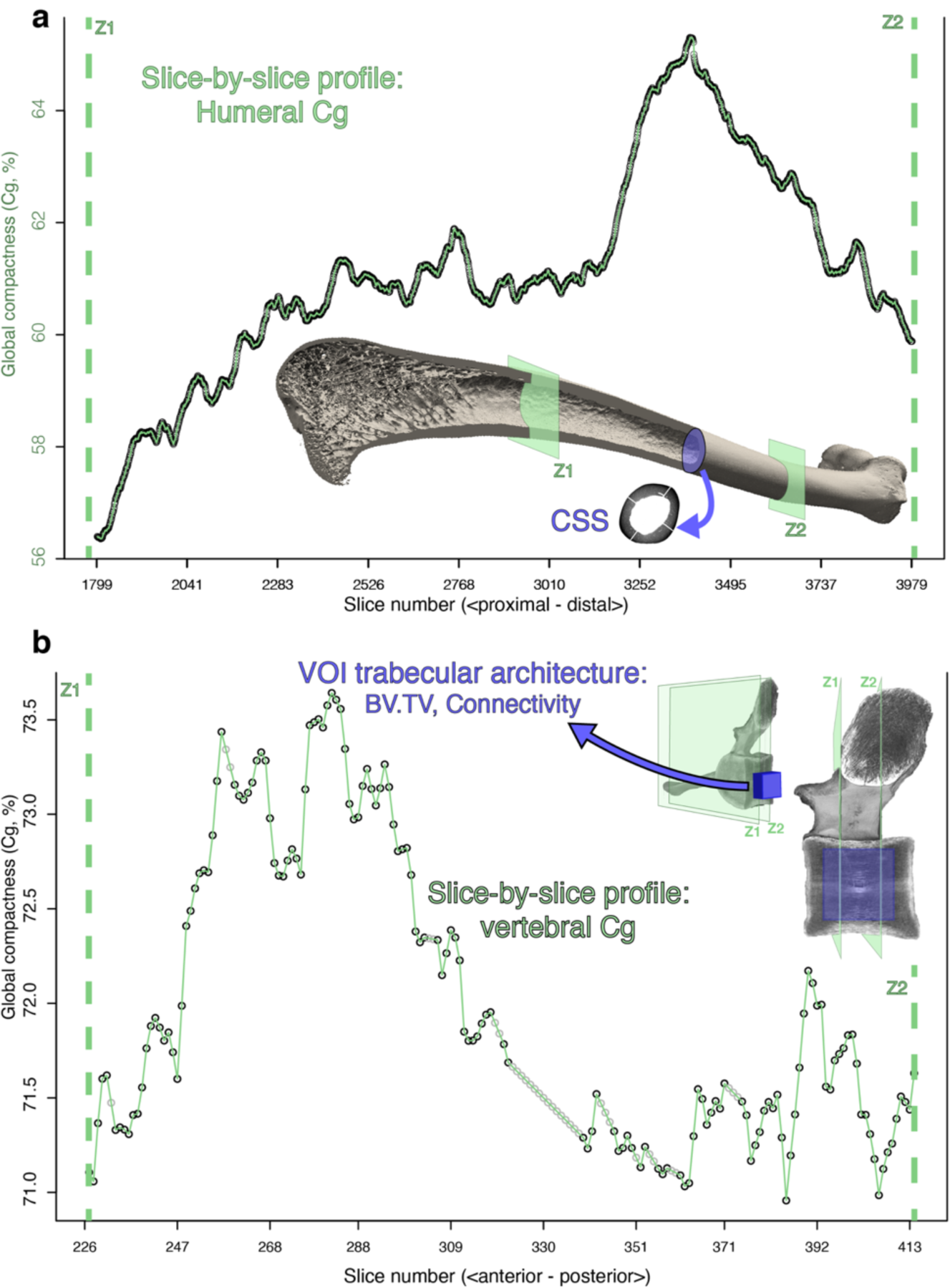
Description of the acquired bone structure parameters. Humeral (**a**) and vertebral (**b**) global compactness (Cg) were acquired along proximodistal and anteroposterior profiles, respectively. For the humerus (**a**), cross-sectional shape (CSS) at midshaft and diaphysis elongation (DE; diaphysis length/total cross-sectional area at midshaft) were also acquired. For the vertebra, a Volume of Interest (VOI) was defined in the centre of the centrum, and the trabecular parameters bone fraction (BV/TV) and Connectivity were acquired. Example bones: humerus of a red fox (*Vulpes vulpes*, ZMB_Mam_49955 and lumbar vertebra of a dugong (*Dugong dugon* ZMB_Mam_69340); not to scale.

Both humerus and lumbar vertebrae were not available for each sampled specimen. Furthermore, some of the CT-scans of Dumont et al. (2013) only captured the centrum, so only parameters of the centrum’s VOI could be acquired. Therefore, the analyses were run with three datasets: whole humerus (159 specimens), whole vertebra (160 specimens), and vertebral centrum (188 specimens). All specialised clades and outgroups were represented in all datasets. Parameters were averaged for those species that were represented by several specimens. All raw values can be found on figshare (doi: 10.6084/m9.figshare.12600440). Descriptive statistics are given in Supplementary Table S2.

In order to avoid potential bias affecting bone structure, specimens were selected to be adult (as indicated by size and/or epiphyseal fusion), devoid of apparent disease, and coming from the wild. But as marsupials and many small-sized mammals maintain unfused humeral epiphyses well into adulthood (if not throughout life), humeral epiphyses ROI parameters were not included in this analysis. Species body mass were taken from the AnAge (Tacutu et al., 2013) and MOM v1.4 (Smith et al., 2003) databases.

### History of lifestyle transitions in mammals

All data analyses were performed with R (R Core Team, 2020) (3.6.2). A reconstruction of the ancestral character states was performed for the tree of extant mammal genera (1167 tips) using stochastic character mapping (make.simmap function, 1000 simulations, equal rate model; phytool package (Revell, 2012)).

### Size effect and lifestyle signal in bone structure parameters

All regressions and AN(C)OVAs were performed with generalised least squares linear models comprising a within-group correlation structure based on the optimised lambda value of the model’s residuals(Revell, 2010) (gls function, nlme package(Pinheiro et al., 2016), corPagel function, APE package; Paradis et al., 2004). Nagelkerke pseudo R^2^ were computed with the rsquared function (piecewiseSEM package; Lefcheck, 2016). For AN(C)OVAs, posthoc pairwise comparisons were performed with the glht and mcp functions (Tukey’s multiple comparisons; multcomp package (Hothorn et al., 2008). Variables were log-transformed when their distribution was blatantly deviating from normality. For Connectivity, for which log-transformation was recommendable, two specimens had a value of 0 (empty VOIs). Their values were hence replaced by the minimum Connectivity otherwise found in the dataset (0.875) subtracted by 10% to make the log-transformation possible.

Specimen size proxies were used to test the effect of body size on the parameters: mean total cross-sectional area (Houssaye and Botton-Divet, 2018) (see above) for the humeral dataset, and centrum length (Houssaye et al., 2019) for the vertebral datasets. In both cases, the correlation with the species body mass was strong (*p*-values < 0.0001; pseudo R^2^ = 0.90 and 0.88, respectively). These traits were hence viewed as satisfactory size proxies. While our sampling endeavoured to cover the whole range of body sizes for each lifestyle, a difference of size was found among them: aquatic taxa are larger than other lifestyles (*p*-values < 0.032; see also Supplementary Methods S2). No conspicuous differences were found between the other lifestyles (*p*-values > 0.129). Furthermore, the terrestrial taxa also feature a greater disparity in their sizes, their range encompassing all other lifestyles but the largest aquatic taxa. Subsets of the datasets for which the terrestrial class content was pruned to match the size and disparity of the other classes were therefore additionally examined: to compare subterranean and aerial lifestyles to the terrestrial one, the latter was pruned to 63 taxa for the humeral dataset (from the 83 original taxa), and to 46 and 48 taxa for the vertebral datasets (from the original 82 and 92 taxa); to compare the aquatic and terrestrial lifestyles, the latter was pruned to 10 taxa for the humeral dataset, and 16 and 24 taxa for the vertebral datasets.

We investigated the effect of size and lifestyle on each studied bone structure parameters using ANCOVAs. When a significant correlation between the size proxy and the parameter under study was recovered, residuals of the corresponding regression were computed. The latter were regarded as the sized-corrected version of these parameters (Revell, 2009), which were used for the convergence analysis, as well as for visualization purposes (phenograms and boxplots, see below).

### Convergence analysis

We quantified the convergent acquisition of each specialised lifestyle using the framework of the convevol package (Stayton, 2015) for R. For each lifestyle, the number and composition of convergent clades was recovered from the result of the stochastic character mapping (Supplementary Fig. S1). While the stochastic character mapping suggested that three genera of petauroids (gliding marsupials and their close relatives) acquired their aerial lifestyle independently, all members of this clade are here classified as non-specialised but non-terrestrial. Because these other petauroids were not sampled (see sampling criteria above), only one convergence was counted for petauroids (see also Supplementary Methods S3). Accordingly, the datasets were aggregated by taking the mean values of each converging clade, as well as each of their respective TSG. The timetree was correspondingly amended by reducing each specialised clade and each sister-group to one tip Supplementary Fig. S2). We used univariate versions of the convevol functions to analyse the evolution of each trait of interest. We computed the convergence index C1 and associated *p*-value (convratsig function), which gives an overall assessment of how much the phenotypes of converging clades evolved towards one another (Stayton, 2015). Because we studied the traits univariately, and because our sampling comprises a non-specialised sister-group for each converging clade, we also investigated the direction of evolution in the last branch leading to each converging clade. We hence wrote a custom function convDir based on convevol’s convnum. For each converging tip, the function assesses whether its value is greater or lower than the reconstructed value of the node directly ancestral to it. The function also assesses whether the tip value falls outside the 95% confidence interval (95CI) of the reconstructed ancestral value. Phenograms (traitgrams) were plotted with the phenogram function (phytools package (Revell, 2012)). Modified convevol functions and convDir are available on GitHub: https://github.com/ https://github.com/eliamson/XXX.

### Evolvability analysis

For traits that were found as converging with the previous analysis, we tested whether the TSG differed from the other terrestrial, more distantly related taxa. For each specialised lifestyle, we used AN(C)OVAs (same methodology as for the size effect and lifestyle signal, see above) comparing three classes: ‘specialised lifestyle’, ‘TSG’, and ‘more distant terrestrial.’ This analysis was not performed for the aquatic clades because both diverged particularly early from their respective sister-groups. Differences were visualised with boxplots.

### Phylogenetic signal

To avoid the potential influence of lifestyles, only terrestrial species were included in this analysis. Pagel’s lambda (phylosig function, phytools package; Revell, 2012) was computed for all terrestrial species taken individually and also after aggregating the TSG (as for the convergence analysis, taking the mean of each clade). This aggregation was used to assess the influence of the most recent diversifications on the phylogenetic signal. The size-corrected trait values (see above) were used when relevant (for all traits but CSS).

## Acknowledgements

We thank A. Bibl, S. Bock, E. Bärmann, C. Funk, M. Herbin, F. Mayer, V. Nicolas, O. Pauwels, R. Portela Miguez, A. Rosemann, G. Veron, D. Willborn, F. Zachos for their help accessing specimens under their care. V. de Buffrénil, B. Clark, V. Fernandez, D. Germain, M. Kirchner, L. Jansen, K. Mahlow, R. Portela Miguez, J. Müller, A. Welter and P. Wils are thanked for helping with the acquisition of CT data. Members of the BMW lab are acknowledged for insightful discussions. We thank T. Stayton for helping with the convevol package. This work was funded by the German Research Council (Deutsche Forschungsgemeinschaft; grant number AM 517/1-1).

## Author contributions

EA contributed to the study design, collected data, run the analyses and drafted the manuscript. FB contributed to the study design and manuscript preparation.

## Data availability

The datasets generated and analysed during the current study are available on figshare (doi: 10.6084/m9.figshare.12600440). Raw CT-scan data are curated by the Museum für Naturkunde (Berlin, Germany) and available upon reasonable request.

## Code availability

Univariate implementation of the functions of the R package convevol (Stayton, 2015), as well as the custom function convDir, are available on GitHub: https://github.com/eliamson/XXX.

## References

Amador LI, Simmons NB, Giannini NP. 2019. Aerodynamic reconstruction of the primitive fossil bat *Onychonycteris finneyi* (Mammalia: Chiroptera). Biol Lett 15:20180857. doi: 10.1098/rsbl.2018.0857

Amson E. 2019. Overall bone structure as assessed by slice-by-slice profile. Evol Biol 46:343–348. doi: 10.1007/s11692-019-09486-6

Amson E, Nyakatura JA. 2018. Palaeobiological inferences based on long bone epiphyseal and diaphyseal structure – the forelimb of xenarthrans (Mammalia). bioRxiv, 318121, ver 5 peer-reviewed Recomm by PCI Paleo. doi: 10.1101/318121

Barak MM, Lieberman DE, Hublin J-J. 2011. A Wolff in sheep’s clothing: Trabecular bone adaptation in response to changes in joint loading orientation. Bone 49:1141–1151. doi: 10.1016/j.bone.2011.08.020

Bardua C, Fabre A, Bon M, Das K, Stanley EL, Blackburn DC, Goswami A. 2020. Evolutionary integration of the frog cranium. Evolution 74:1200–1215. doi: 10.1111/evo.13984

Basu C, Wilson AM, Hutchinson JR. 2019. The locomotor kinematics and ground reaction forces of walking giraffes. J Exp Biol 222:jeb159277. doi: 10.1242/jeb.159277

Beard KC. 1993. Origin and Evolution of Gliding in Early Cenozoic Dermoptera (Mammalia, Primatomorpha)Primates and Their Relatives in Phylogenetic Perspective. Boston, MA: Springer US. pp. 63–90. doi: 10.1007/978-1-4899-2388-2_2

Biewener AA. 2005. Biomechanical consequences of scaling. J Exp Biol 208:1665–1676. doi: 10.1242/jeb.01520

Blomberg SP, Garland T, Ives AR. 2003. Testing for phylogenetic signal in comparative data: behavioral traits are more labile. Evolution 57:717–745.

Briggs DEG. 2017. Seilacher, Konstruktions-Morphologie, morphodynamics, and the evolution of form. J Exp Zool Part B Mol Dev Evol 328:197–206. doi: 10.1002/jez.b.22725

Buchholtz EA. 2001. Vertebral osteology and swimming style in living and fossil whales (Order: Cetacea). J Zool 253:175–190. doi: 10.1017/S0952836901000164

Buffrénil V de, Canoville A, D’Anastasio R, Domning DP. 2010. Evolution of sirenian pachyosteosclerosis, a model-case for the study of bone structure in aquatic tetrapods. J Mamm Evol 17:101–120. doi: 10.1007/s10914-010-9130-1

Campione NE, Evans DC. 2012. A universal scaling relationship between body mass and proximal limb bone dimensions in quadrupedal terrestrial tetrapods. BMC Biol 10:60. doi: 10.1186/1741-7007-10-60

Canoville A, Laurin M. 2010. Evolution of humeral microanatomy and lifestyle in amniotes, and some comments on palaeobiological inferences. Biol J Linn Soc 100:384–406. doi: 10.1111/j.1095-8312.2010.01431.x

Chirchir H, Kivell TL, Ruff CB, Hublin J-J, Carlson KJ, Zipfel B. 2015. Recent origin of low trabecular bone density in modern humans. Proc Natl Acad Sci 112:366–371. doi: 10.1073/pnas.1411696112

Cubo J, Legendre P, de Ricqlès A, Montes L, de Margerie E, Castanet J, Desdevises Y. 2008. Phylogenetic, functional, and structural components of variation in bone growth rate of amniotes. Evol Dev 10:217–27. doi: 10.1111/j.1525-142X.2008.00229.x

Currey JD. 2003. The many adaptations of bone. J Biomech 36:1487–1495. doi: 10.1016/S0021-9290(03)00124-6

Currey JD, Alexander RM. 1985. The thickness of the walls of tubular bones. J Zool 206:453–468. doi: 10.1111/j.1469-7998.1985.tb03551.x

Domning DP. 1976. An ecological model for late Tertiary sirenian evolution in the North Pacific ocean. Syst Zool 25:352. doi: 10.2307/2412510

Doube M, Kłosowski MM, Arganda-Carreras I, Cordelières FP, Dougherty RP, Jackson JS, Schmid B, Hutchinson JR, Shefelbine SJ. 2010. BoneJ: Free and extensible bone image analysis in ImageJ. Bone 47:1076–1079. doi: 10.1016/j.bone.2010.08.023

Dumont ER. 2010. Bone density and the lightweight skeletons of birds. Proc Biol Sci 277:2193–8. doi: 10.1098/rspb.2010.0117

Dumont M, Laurin M, Jacques F, Pellé E, Dabin W, Buffrénil V de. 2013. Inner architecture of vertebral centra in terrestrial and aquatic mammals: A two-dimensional comparative study. J Morphol 274:570–584. doi: 10.1002/jmor.20122

Ellerman JR. 1956. The subterranean mammals of the world. Trans R Soc South Africa 35:11–20. doi: 10.1080/00359195609519005

Eswaran SK, Gupta A, Adams MF, Keaveny TM. 2005. Cortical and trabecular load sharing in the human vertebral body. J Bone Miner Res 21:307–314. doi: 10.1359/jbmr.2006.21.2.307

Fabre A-C, Cornette R, Goswami A, Peigné S. 2015. Do constraints associated with the locomotor habitat drive the evolution of forelimb shape? A case study in musteloid carnivorans. J Anat 226:596–610. doi: 10.1111/joa.12315

Gasc JP, Jouffroy FK, Renous S. 1986. Morphofunctional study of the digging system of the Namib Desert Golden mole (*Eremitalpa granti namibensis*): cinefluorographical and anatomical analysis. J Zool 208:9–35.

Grossnickle DM, Smith SM, Wilson GP. 2019. Untangling the Multiple Ecological Radiations of Early Mammals. Trends Ecol Evol 11:1–14. doi: 10.1016/j.tree.2019.05.008

Guo X., Kim C. 2002. Mechanical consequence of trabecular bone loss and its treatment: a three-dimensional model simulation. Bone 30:404–411. doi: 10.1016/S8756-3282(01)00673-1

Habib MB, Ruff CB. 2008. The effects of locomotion on the structural characteristics of avian limb bones. Zool J Linn Soc 153:601–624. doi: 10.1111/j.1096-3642.2008.00402.x

Halpert AP, Jenkins FA, Franks H. 1987. Structure and scaling of the lumbar vertebrae in African bovids (Mammalia: Artiodactyla). J Zool 211:239–258. doi: 10.1111/j.1469-7998.1987.tb08599.x

Hedrick BP, Dickson B V., Dumont ER, Pierce SE. 2020. The evolutionary diversity of locomotor innovation in rodents is not linked to proximal limb morphology. Sci Rep 10:717. doi: 10.1038/s41598-019-57144-w

Hothorn T, Bretz F, Westfall P. 2008. Simultaneous inference in general parametric models. Biometrical J 50:346–363. doi: 10.1002/bimj.200810425

Houssaye A, Boistel R, Böhme W, Herrel A. 2013. Jack-of-all-trades master of all? Snake vertebrae have a generalist inner organization. Naturwissenschaften 100:997–1006. doi: 10.1007/s00114-013-1102-x

Houssaye A, Botton-Divet L. 2018. From land to water: evolutionary changes in long bone microanatomy of otters (Mammalia: Mustelidae). Biol J Linn Soc. doi: 10.1093/biolinnean/bly118

Houssaye A, Herrel A, Boistel R, Rage J. 2019. Adaptation of the vertebral inner structure to an aquatic life in snakes: Pachyophiid peculiarities in comparison to extant and extinct forms. Comptes Rendus Palevol 18:783–799. doi: 10.1016/j.crpv.2019.05.004

Houssaye A, Martin Sander P, Klein N. 2016. Adaptive patterns in aquatic amniote bone microanatomy—more complex than previously thought. Integr Comp Biol 56:1349–1369. doi: 10.1093/icb/icw120

Houssaye A, Mazurier A, Herrel A, Volpato V, Tafforeau P, Boistel R, Buffrénil V de. 2010. Vertebral microanatomy in squamates: structure, growth and ecological correlates. J Anat 217:715–27. doi: 10.1111/j.1469-7580.2010.01307.x

Houssaye A, Tafforeau P, Herrel A. 2014. Amniote vertebral microanatomy–what are the major trends? Biol J Linn Soc 112:735–746.

Howell AB. 1930. Aquatic mammals: their adaptations to life in the water. Springfield: Charles C. Thomas.

Hu Y, Albertson RC. 2016. Developmental Biases on Morphological EvolvabilityEncyclopedia of Evolutionary Biology. pp. 399–403. doi: 10.1016/B978-0-12-800049-6.00134-7

Jackson SM, Thorington RW. 2012. Gliding mammals: Taxonomy of living and extinct species. Smithson Contrib to Zool 638:1–117. doi: 10.5479/si.00810282.638.1

Jones KE. 2015. Evolutionary allometry of the thoracolumbar centra in felids and bovids. J Morphol 276:818–831. doi: 10.1002/jmor.20382

Jones KE, Pierce SE. 2016. Axial allometry in a neutrally buoyant environment: effects of the terrestrial-aquatic transition on vertebral scaling. J Evol Biol 29:594–601. doi: 10.1111/jeb.12809

Kilbourne BM, Hutchinson JR. 2019. Morphological diversification of biomechanical traits: Mustelid locomotor specializations and the macroevolution of long bone cross-sectional morphology. BMC Evol Biol 19:1–16. doi: 10.1186/s12862-019-1349-8

Kivell TL. 2016. A review of trabecular bone functional adaptation: what have we learned from trabecular analyses in extant hominoids and what can we apply to fossils? J Anat 228:569–594. doi: 10.1111/joa.12446

Kley NJ, Kearney M. 2007. Adaptations for digging and burrowing In: Hall BK, editor. Fins into Limbs: Evolution, Development, and Transformation. Chicago: University of Chicago Press. pp. 284–309.

Kojeszewski T, Fish FE. 2007. Swimming kinematics of the Florida manatee (*Trichechus manatus latirostris*): hydrodynamic analysis of an undulatory mammalian swimmer. J Exp Biol 210:2411–8. doi: 10.1242/jeb.02790

Kumar S, Stecher G, Suleski M, Hedges SB. 2017. TimeTree: A resource for timelines, timetrees, and divergence times. Mol Biol Evol 34:1812–1819. doi: 10.1093/molbev/msx116

Lanyon L, Rubin C. 1985. Functional adaptation in skeletal structures In: Hildebrand M, editor. Functional Vertebrate Morphology. Cambridge: Cambridge University Press. pp. 1–25.

Lefcheck JS. 2016. piecewiseSEM: Piecewise structural equation modelling in R for ecology, evolution, and systematics. Methods Ecol Evol 7:573–579. doi: 10.1111/2041-210X.12512

Lister AM. 2014. Behavioural leads in evolution: evidence from the fossil record. Biol J Linn Soc 112:315–331. doi: 10.1111/bij.12173

MacLarnon A. 1996. The scaling of gross dimensions of the spinal cord in primates and other species. J Hum Evol 30:71–87. doi: 10.1006/jhev.1996.0005

Meier PS, Bickelmann C, Scheyer TM, Koyabu D, Sánchez-Villagra MR. 2013. Evolution of bone compactness in extant and extinct moles (Talpidae): exploring humeral microstructure in small fossorial mammals. BMC Evol Biol 13:55. doi: 10.1186/1471-2148-13-55

Mein P, Romaggi JP. 1991. Un gliridé (Mammalia, Rodentia) planeur dans le miocène supérieur de l’ardèche: Une adaptation non retrouvée dans la nature actuelle. Geobios 13:45–50. doi: 10.1016/S0016-6995(66)80008-6

Mittra E, Rubin C, Qin YX. 2005. Interrelationship of trabecular mechanical and microstructural properties in sheep trabecular bone. J Biomech 38:1229–1237. doi: 10.1016/j.jbiomech.2004.06.007

Navalón G, Bright JA, Marugán-Lobón J, Rayfield EJ. 2019. The evolutionary relationship among beak shape, mechanical advantage, and feeding ecology in modern birds*. Evolution 73:422–435. doi: 10.1111/evo.13655

Nazarian A, Von Stechow D, Zurakowski D, Müller R, Snyder BD. 2008. Bone volume fraction explains the variation in strength and stiffness of cancellous bone affected by metastatic cancer and osteoporosis. Calcif Tissue Int 83:368–379. doi: 10.1007/s00223-008-9174-x

Nowak RM. 1999. Walker’s Mammals of the World, Walker’s Mammals of the World. Baltimore, MD: John Hopkins University Press. doi: 10.1353/book.59141

Paradis E, Claude J, Strimmer K. 2004. APE: Analyses of phylogenetics and evolution in R language. Bioinformatics 20:289–290. doi: 10.1093/bioinformatics/btg412

Parins-Fukuchi C. 2020. Mosaic evolution, preadaptation, and the evolution of evolvability in apes. Evolution 74:297–310. doi: 10.1111/evo.13923

Patel BA, Ruff CB, Simons ELR, Organ JM. 2013. Humeral cross-sectional shape in suspensory primates and sloths. Anat Rec 556:545–556. doi: 10.1002/ar.22669

Pinheiro J, Bates D, DebRoy S, Sarkar D, R Core Team. 2016. nlme: Linear and Nonlinear Mixed Effects Models. R package version 3.1-128.

R Core Team. 2020. R: A language and environment for statistical computing.

Raia P, Carotenuto F, Meloro C, Piras P, Pushkina D. 2010. The shape of contention: Adaptation, history, and contingency in ungulate mandibles. Evolution 64:1489–1503. doi: 10.1111/j.1558-5646.2009.00921.x

Revell LJ. 2012. phytools: An R package for phylogenetic comparative biology (and other things). Methods Ecol Evol 3:217–223. doi: 10.1111/j.2041-210X.2011.00169.x

Revell LJ. 2010. Phylogenetic signal and linear regression on species data. Methods Ecol Evol 1:319–329. doi: 10.1111/j.2041-210X.2010.00044.x

Revell LJ. 2009. Size-correction and principal components for interspecific comparative studies. Evolution 63:3258–68. doi: 10.1111/j.1558-5646.2009.00804.x

Ricqlès A de, Buffrénil V de. 2001. Bone histology, heterochronies and the return of tetrapods to life in water: were are we In: Mazin J, Buffrénil V, editors. Secondary Adaptation of Tetrapods to Life in Water. München: Verlag Dr Friedrich Pfeil. pp. 289–310.

Ruff CB, Hayes WC. 1983. Cross-sectional geometry of Pecos Pueblo femora and tibiae – a biomechanical investigation: I. Method and general patterns of variation. Am J Phys Anthropol 60:359–81. doi: 10.1002/ajpa.1330600308

Runestad JA, Ruff CB. 1995. Structural adaptations for gliding in mammals with implications for locomotor behavior in paromomyids. Am J Phys Anthropol 98:101–119. doi: 10.1002/ajpa.1330980202

Schindelin J, Arganda-Carreras I, Frise E, Kaynig V, Longair M, Pietzsch T, Preibisch S, Rueden C, Saalfeld S, Schmid B, Tinevez J-Y, White DJ, Hartenstein V, Eliceiri K, Tomancak P, Cardona A. 2012. Fiji: an open-source platform for biological-image analysis. Nat Methods 9:676–682. doi: 10.1038/nmeth.2019

Seilacher A. 1970. Arbeitskonzept zur Konstruktions-Morphologie. Lethaia 3:393–396.

Shefelbine SJ, Tardieu C, Carter DR. 2002. Development of the femoral bicondylar angle in hominid bipedalism. Bone 30:765–770. doi: 10.1016/S8756-3282(02)00700-7

Slijper EJ. 1946. Comparative biologic-anatomical investigations on the vertebral column and spinal musculature of mammals. Verh der K Ned Akad van Wet Natuurkd 42:1–128.

Smith FA, Lyons SK, Ernest SKM, Jones KE, Kaufman DM, Dayan T, Marquet PA, Brown JH, Haskell JP. 2003. Body mass of late Quaternary mammals. Ecology 84:3403. doi: 10.1890/02-9003

Smith SM, Angielczyk KD. 2020. Deciphering an extreme morphology: bone microarchitecture of the hero shrew backbone (Soricidae: *Scutisorex*). Proc R Soc B Biol Sci 287:20200457. doi: 10.1098/rspb.2020.0457

Sode M, Burghardt AJ, Nissenson RA, Majumdar S. 2008. Resolution dependence of the monmetric trabecular structure indices. Bone 42:728–736. doi: 10.1016/j.bone.2007.12.004

Stanchak KE, Arbour JH, Santana SE. 2019. Anatomical diversification of a skeletal novelty in bat feet. Evolution 1–13. doi: 10.1111/evo.13786

Stayton CT. 2015. The definition, recognition, and interpretation of convergent evolution, and two new measures for quantifying and assessing the significance of convergence. Evolution 69:2140–2153. doi: 10.1111/evo.12729

Storch G, Engesser B, Wuttke M. 1996. Oldest fossil record of gliding in rodents. Nature 379:439–441. doi: 10.1038/379439a0

Swartz SM, Bennett MB, Carrier DR. 1992. Wing bone stresses in free flying bats and the evolution of skeletal design for flight. Nature 359:726–729. doi: 10.1038/359726a0

Tacutu R, Craig CT, Budovsky A, Wuttke D, Lehmann G, Taranukha D, Costa J, Fraifeld VE, D. Magalhães J. 2013. Human Ageing Genomic Resources: Integrated databases and tools for the biology and genetics of ageing. Nucleic Acids Res 41:D1027–D1033. doi: 10.1093/nar/gks1155

Voeten DFAE, Cubo J, de Margerie E, Röper M, Beyrand V, Bureš S, Tafforeau P, Sanchez S. 2018. Wing bone geometry reveals active flight in *Archaeopteryx*. Nat Commun 9:923. doi: 10.1038/s41467-018-03296-8

