## Supplementary for "Differing effects of size and lifestyle on bone structure in mammals"

***Supplementary Methods S1. TimeTree alterations.*** **2**
***Supplementary Methods S2. Body size ~ lifestyle.*** **3**
***Supplementary Methods S3. Petauroid lifestyle.*** **4**
***Supplementary Figure S1. Timetree of sampled.*** **5**
***Supplementary Figure S2. Timetree of converging clades.*** **6**
***Supplementary Table S1. Specimen list.*** **7**
***Supplementary Table S2. Descriptive statistics*** **12**
***Supplementary Results S1. AN(C)OVAs*** **13**
***Supplementary Results S2. Phenograms.*** **19**
***Supplementary Results S3. Phylogenetic signal.*** **25**
***References*** **28**

**Supplementary Methods S1. TimeTree alterations.**

Alterations brought to the TimeTree of Kumar et al.(Kumar et al. 2017).

Two species names were corrected:

*Aotus azarai* => *Aotus azarae* (according to Wilson & Reeder (Wilson & Reeder 2005)).

*Tatera* sp. KIK1704 => *Tatera indica*

Seven species names were swapped with other species of the same genus (straightforward renaming as only one species of each of these genera was sampled):

*Rhizomys pruinosus* => *Rhizomys sumatrensis*

*Paraechinus aethiopicus* => *Paraechinus hypomelas*

*Petinomys setosus* => *Petinomys fuscocapilus*

*Abrocoma cinerea* => *Abrocoma budini*

*Hylomys parvus* => *Hylomys megalotis*

*Ctenomys torquatus* => *Ctenomys brasiliensis*

*Petaurillus kinlochii* => *Petaurillus hosei*

Four species were added to the tree using external sources:

*Anomalurus pelii*: divergence from most closely related sampled species (*A. derbianus*) at 12.2 Ma (Fabre et al. 2018)

*Idiurus macrotis*: divergence from *I. zenkeri* at 11.2 Ma (P.-H. Fabre, pers. comm.)

*Oryzorictes tetradactylus*: divergence from most closely related sampled species (*O. hova*) at 5.13 Ma (Everson et al. 2018)

*Hydrodamalis gigas*: divergence from most closely related sampled species (*Dugong dugon*) at 28.59 Ma (Springer et al. 2015).

The final, altered tree (182 species) can be found on figshare (doi: 10.6084/m9.figshare.12600440).

**Supplementary Methods S2. Body size ~ lifestyle.**
Distribution of body size according to lifestyle.

The log-transformed vertebral centrum length is used here as a size proxy. Abbreviations: Ae, Aerial; Aq, Aquatic; Su, Subterreanean; Te, terrestrial.

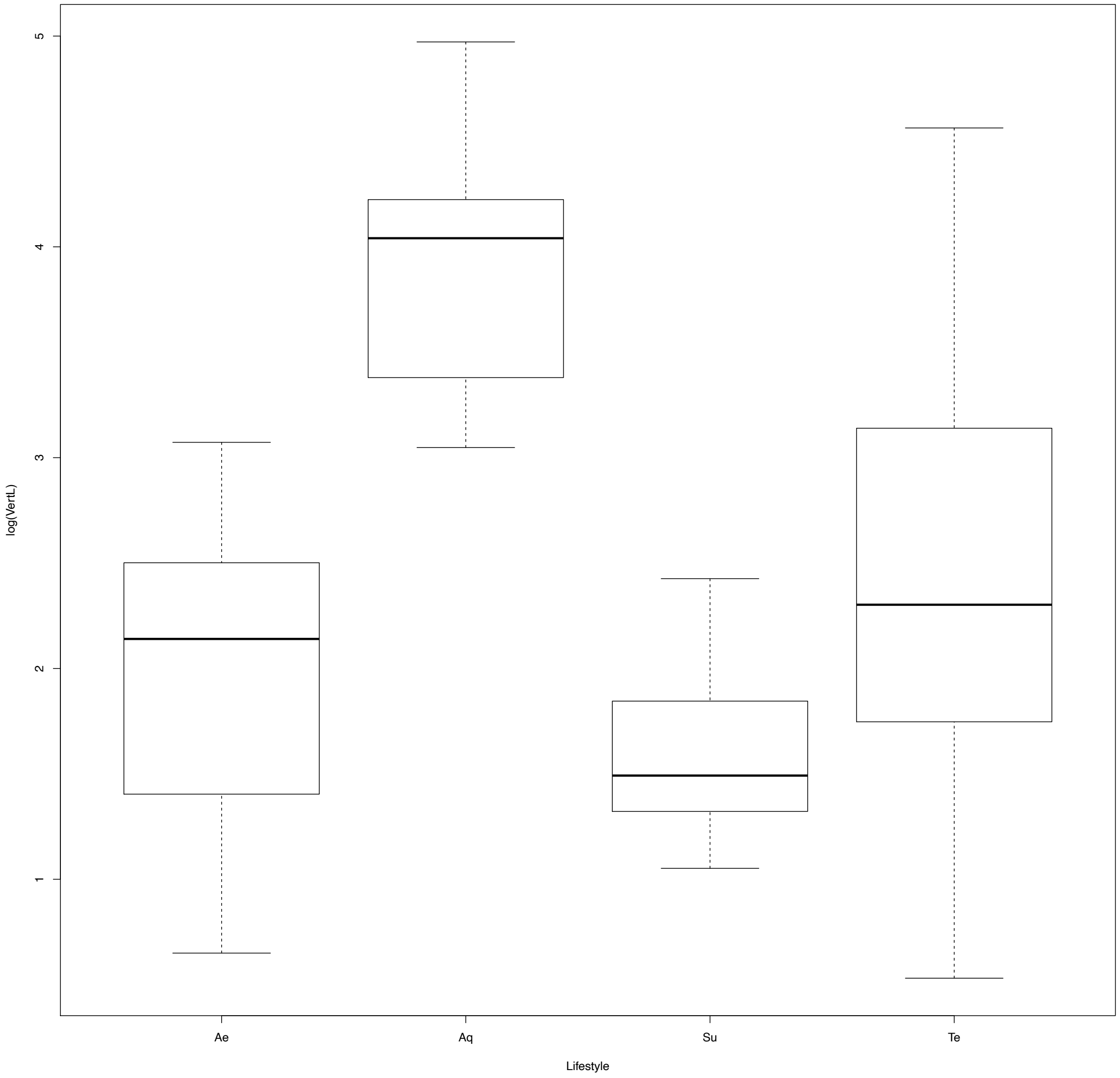


Result of post-hoc pairwise comparisons (glht function, multcomp package (Hothorn et al. 2008):

| | Estimate | Std. Error | z value | Pr(>\|z\|) | |
|---|---|---|---|---|---|
| Aq - Ae == 0 | 1.7962 | 0.5149 | 3.488 | 0.00234 | ** |
| Su - Ae == 0 | 0.3855 | 0.3224 | 1.196 | 0.60019 | |
| Te - Ae == 0 | 0.6303 | 0.2973 | 2.12 | 0.12903 | |
| Su - Aq == 0 | -1.4106 | 0.4546 | -3.103 | 0.00862 | ** |
| Te - Aq == 0 | -1.1658 | 0.4343 | -2.684 | 0.03118 | * |
| Te - Su == 0 | 0.2448 | 0.15 | 1.632 | 0.33052 | |

### **Supplementary Methods S3. Petauroid lifestyle.**

Acquisitions of gliding lifestyle in Petauroidea.

The most probable reconstructed ancestral state for Petauroidea is terrestrial (species tree from TimeTree.org (Kumar et al. 2017); make.simmap function of the phytools package (Revell 2012), model with equal rates of transition, 1000 simulations), implying that an aerial lifestyle was convergently acquired in *Acrobates*, *Petauroides*, and *Petaurus*. However, all other petauroids are arboreal/scansorial (Nowak 2020). The latter were hence not sampled (see Methods of main text), and only one acquisition of the aerial lifestyle was counted for Petauroidea.

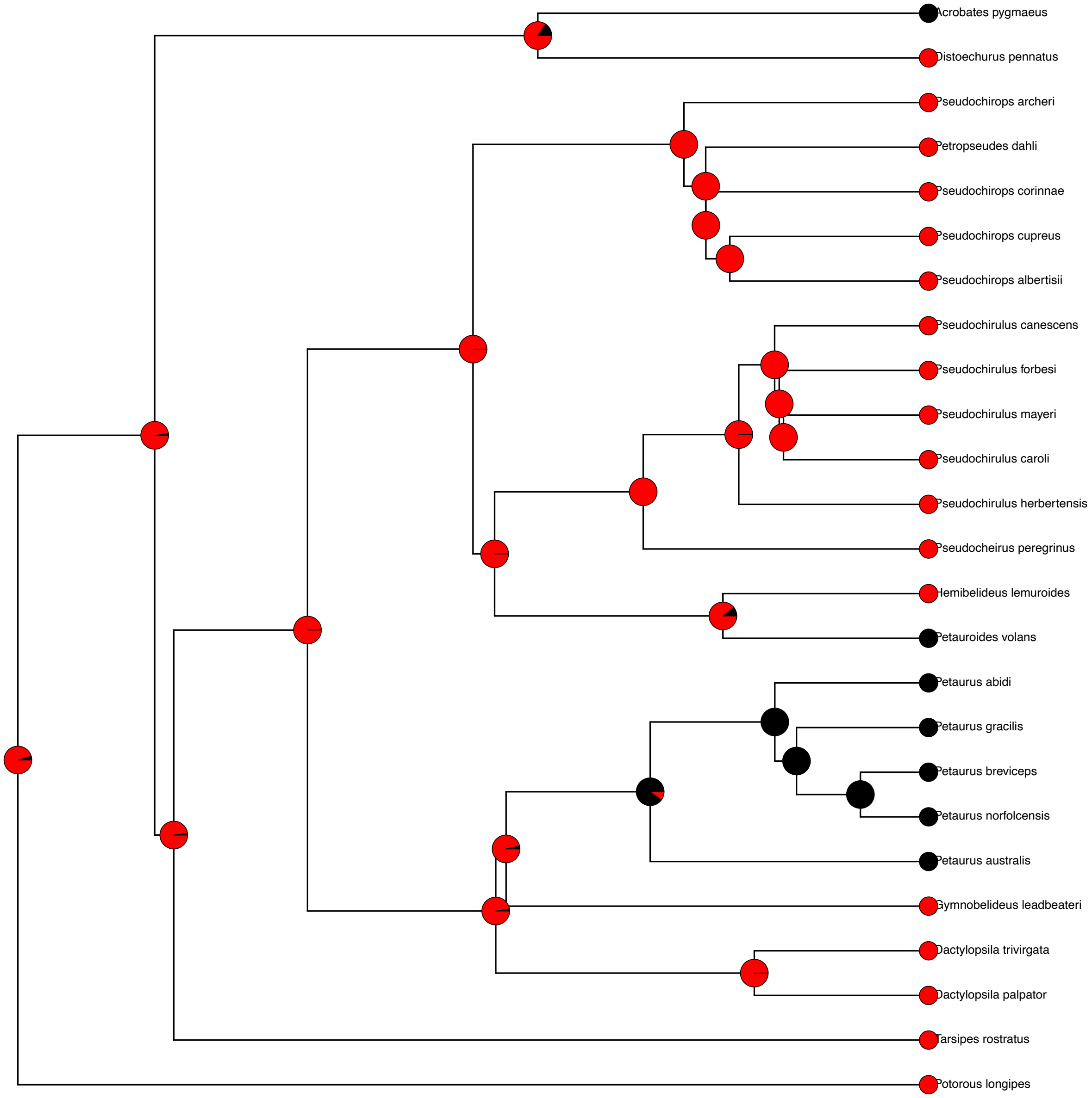

**Supplementary Figure S1. Timetree of sampled.**

Timetree of the mammalian species sampled.

Colours correspond to the specialised lifestyles, i.e., aerial, aquatic, and subterranean; black corresponds to the terrestrial lifestyle. States at the nodes are reconstructed with stochastic character mapping (make.simmap function, 1000 simulations, equal rate model; phytool(Revell 2012). Each doodle represents an independent acquisition of one of the three specialised lifestyle. Abbreviations: Anom., Anomaluromorpha; Lag., Lagomorpha; Xe, Xenarthra.

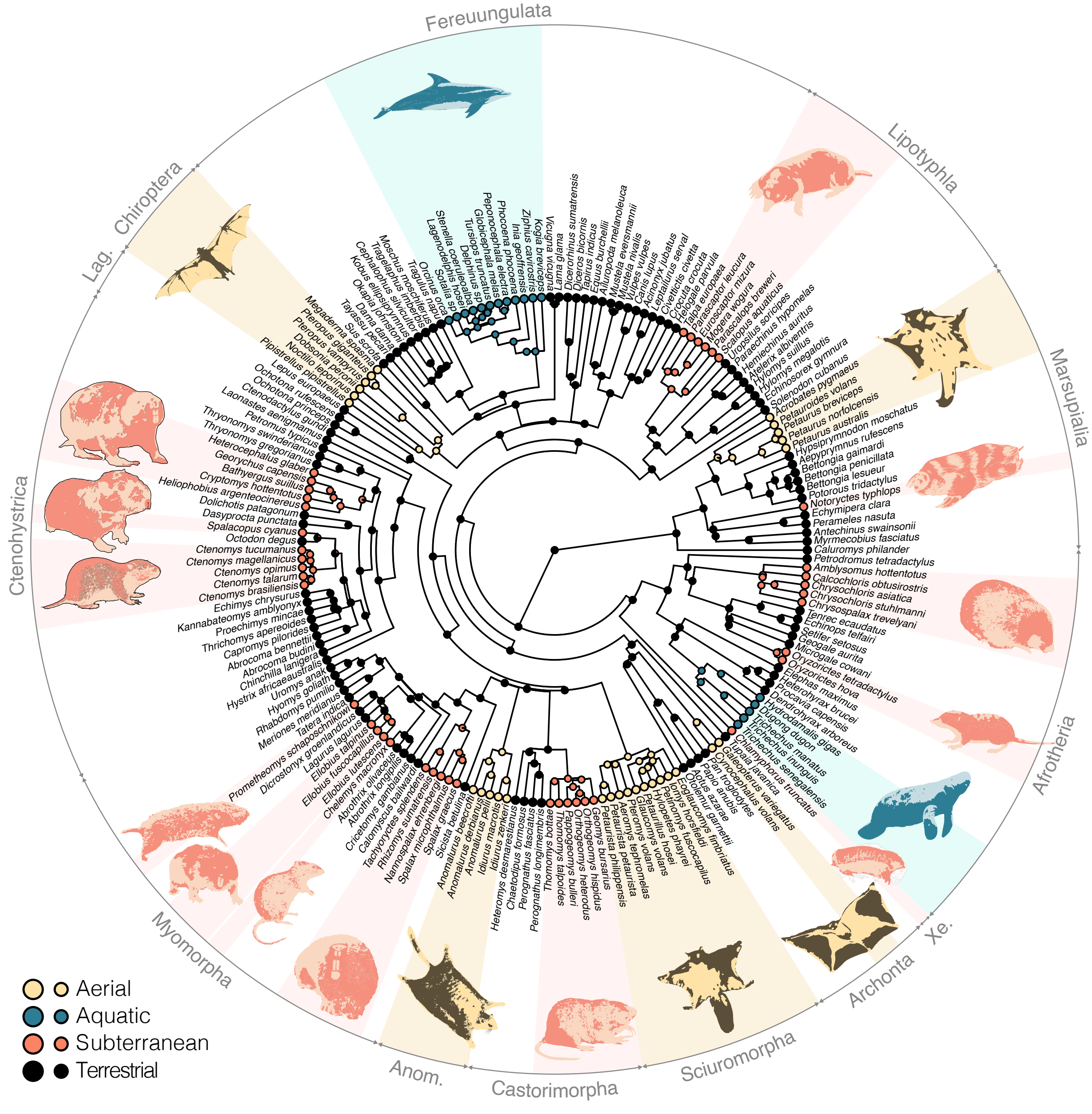

### Supplementary Figure S2. Timetree of converging clades.

Timetree amended by reducing each specialised clade and each sister-clade to one tip.

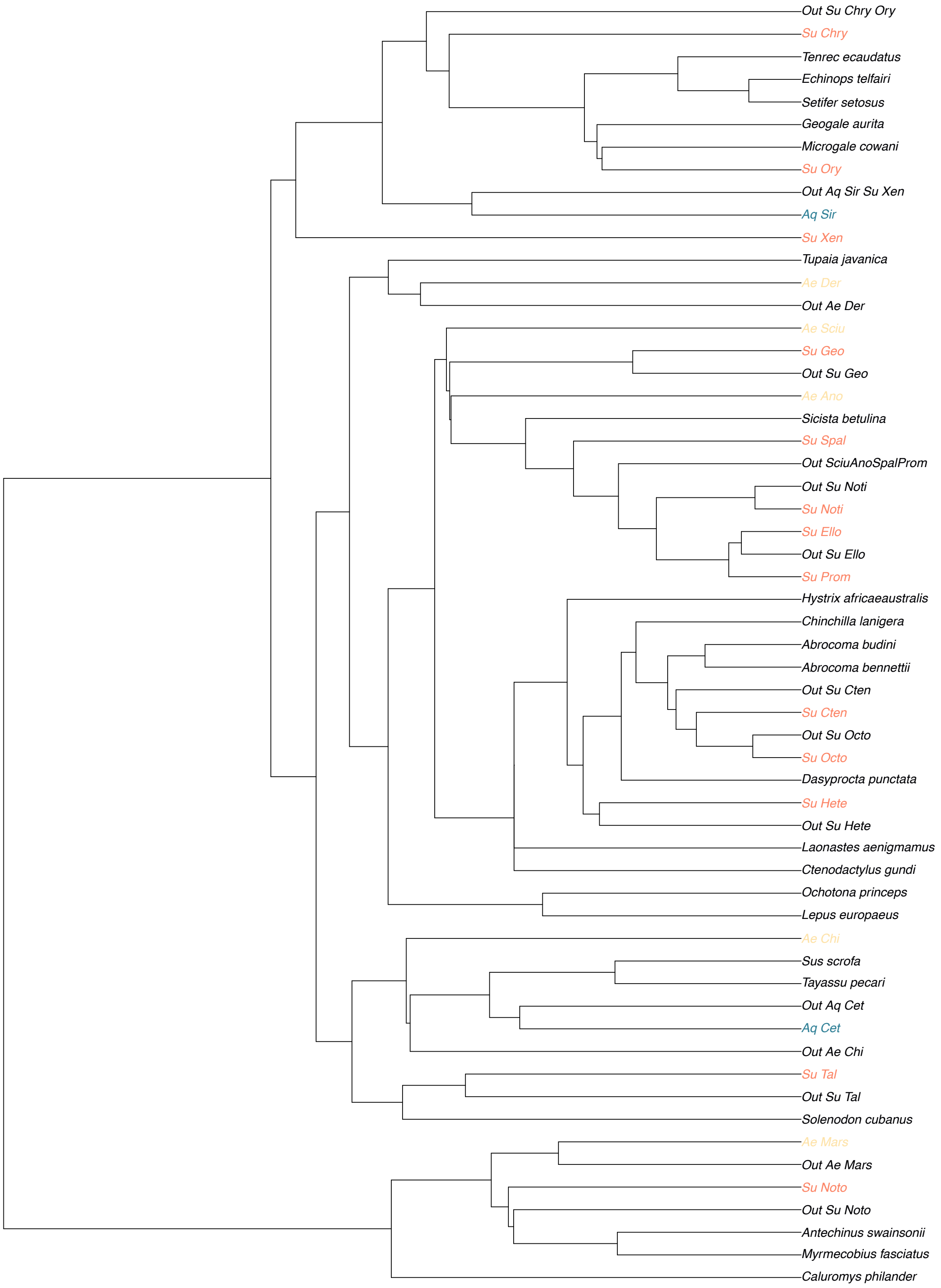

### Supplementary Table S1. Specimen list.

List of all sampled specimens, with their corresponding lifestyle and abbreviated name of their specialised clade, terrestrial sister-group of a specialised clade, or more distantly related terrestrial clade (the latter are referred to as "Else"). See corresponding timetree as Supplementary Figure 2.

| Species | Collection number | Lifestyle | Clade abbreviation |
|---|---|---|---|
| *Anomalurus beecrofti* | ZMB_Mam_36475 | Ae | Ae_Ano |
| *Anomalurus derbianus* | ZMB_Mam_36344 | Ae | Ae_Ano |
| *Anomalurus pelii* | ZMB_Mam_18404 | Ae | Ae_Ano |
| *Idiurus macrotis* | ZMB_Mam_10088 | Ae | Ae_Ano |
| *Idiurus zenkeri* | ZMB_Mam_22747 | Ae | Ae_Ano |
| *Dobsonia peronii* | ZMB_Mam_66825 | Ae | Ae_Chi |
| *Megaderma spasma* | ZMB_Mam_85644 | Ae | Ae_Chi |
| *Noctilio leporinus* | ZMB_Mam_85657 | Ae | Ae_Chi |
| *Pipistrellus pipistrellus* | ZMB_Mam_55128 | Ae | Ae_Chi |
| *Pteropus giganteus* | MNHN_A_1251 | Ae | Ae_Chi |
| *Pteropus vampyrus* | ZMB_Mam_88424 | Ae | Ae_Chi |
| *Galeopterus variegatus* | ZMB_Mam_69096 | Ae | Ae_Der |
| *Cynocephalus volans* | ZMB_Mam_69105 | Ae | Ae_Der |
| *Acrobates pygmaeus* | ZMB_Mam_6241 | Ae | Ae_Mars |
| *Petauroides volans* | ZMB_Mam_106893 | Ae | Ae_Mars |
| *Petaurus australis* | MNHN-ZM-MO1883-1533 | Ae | Ae_Mars |
| *Petaurus breviceps* | ZMB_Mam_104697 | Ae | Ae_Mars |
| *Petaurus norfolcensis* | MNHN-ZM-MO1961-951 | Ae | Ae_Mars |
| *Aeromys tephromelas* | NHMUK 1977.2886 | Ae | Ae_Sciu |
| *Eoglaucomys fimbriatus* | NHMUK GERM 1721.a | Ae | Ae_Sciu |
| *Glaucomys volans* | MNHN_ZM_MO_1951_1023 | Ae | Ae_Sciu |
| *Hylopetes phayrei* | NHMUK ZD 1994.199 | Ae | Ae_Sciu |
| *Iomys horsfieldi* | NHMUK 1977.2885 | Ae | Ae_Sciu |
| *Petaurillus hosei* | NHMUK ZD 1900.7.29.26 | Ae | Ae_Sciu |
| *Petaurista petaurista* | ZMB_Mam_78560 | Ae | Ae_Sciu |
| *Petinomys fuscocapilus* | NHMUK 1977.465 | Ae | Ae_Sciu |
| *Petaurista philippensis* | ZMB_Mam_91292 | Ae | Ae_Sciu |
| *Pteromys volans* | ZMB_Mam_78602 | Ae | Ae_Sciu |
| *Delphinus sp* | ZMB_Mam_697.59 | Aq | Aq_Cet |
| *Globicephala melas* | MNHN_LR_M_961 | Aq | Aq_Cet |
| *Inia geoffrensis* | ZMB_Mam_41500 | Aq | Aq_Cet |
| *Kogia breviceps* | MNHN_LR_M_1644 | Aq | Aq_Cet |
| *Lagenodelphis hosei* | MNHN_LR_M1687 | Aq | Aq_Cet |
| *Orcinus orca* | MNHN_1880_260 | Aq | Aq_Cet |
| *Peponocephala electra* | MNHN_LR_M_103.08.076 | Aq | Aq_Cet |
| *Phocoena phocoena* | MNHN_LR_M_958 | Aq | Aq_Cet |
| *Sotalia sp* | ZMB_Mam_35828 | Aq | Aq_Cet |
| *Stenella coeruleoalba* | MNHN_LR_M1848 | Aq | Aq_Cet |
| *Tursiops truncatus* | MNHN_LR_M_1127 | Aq | Aq_Cet |
| *Ziphius cavirostris* | MNHN_LR_M_942 | Aq | Aq_Cet |
| *Dugong dugon* | ZMB_Mam_69340 | Aq | Aq_Sir |
| *Hydrodamalis gigas* | MNHN_AC_1919-48 | Aq | Aq_Sir |
| *Trichechus manatus* | ZMB_Mam_17377 | Aq | Aq_Sir |

| | | | |
|---|---|---|---|
| *Trichechus senegalensis* | ZMB_Mam_69334 | Aq | Aq_Sir |
| *Trichechus inunguis* | ZMB_Mam_35805 | Aq | Aq_Sir |
| *Amblysomus hottentotus* | NMW26095 | Su | Su_Chry |
| *Amblysomus hottentotus* | NMW7194 | Su | Su_Chry |
| *Calcochloris obtusirostris* | ZMB_Mam_35173 | Su | Su_Chry |
| *Chrysochloris stuhlmanni* | NHMUK 1934.4.1.8 | Su | Su_Chry |
| *Chrysochloris asiatica* | ZMB_Mam_76897 | Su | Su_Chry |
| *Chrysochloris asiatica* | NMW970 | Su | Su_Chry |
| *Chrysospalax trevelyani* | NHMUK ZE 1951.11.14.4 | Su | Su_Chry |
| *Ctenomys brasiliensis* | NHMUK ZD 1897.10.3.68 | Su | Su_Cten |
| *Ctenomys opimus* | IRSNB 13028 | Su | Su_Cten |
| *Ctenomys talarum* | NHMUK ZE 1968.8.20.1 | Su | Su_Cten |
| *Ctenomys tucumanus* | ZMB_Mam_18180 | Su | Su_Cten |
| *Ctenomys magellanicus* | ZMB_Mam_38821 | Su | Su_Cten |
| *Ellobius fuscocapillus* | NMW12052 | Su | Su_Ello |
| *Ellobius lutescens* | NMW20329 | Su | Su_Ello |
| *Ellobius talpinus* | ZMB_Mam_78541 | Su | Su_Ello |
| *Orthogeomys heterodus* | ZMB_Mam_106984 | Su | Su_Geo |
| *Orthogeomys hispidus* | NHMUK ZD 1993.120 | Su | Su_Geo |
| *Pappogeomys bulleri* | NHMUK 93.2.5.45 | Su | Su_Geo |
| *Thomomys bottae* | ZMB_Mam_078534 | Su | Su_Geo |
| *Thomomys talpoides* | NMW63141 | Su | Su_Geo |
| *Thomomys talpoides* | NMW63144 | Su | Su_Geo |
| *Bathyergus suillus* | ZMB_Mam_107241 | Su | Su_Hete |
| *Cryptomys hottentotus* | NMW18413 | Su | Su_Hete |
| *Geomys bursarius* | NHMUK 1384.a | Su | Su_Hete |
| *Georychus capensis* | NHMUK ZD 2019.218 | Su | Su_Hete |
| *Georychus capensis* | ZMB_Mam_106888 | Su | Su_Hete |
| *Heterocephalus glaber* | ZMB_Mam_79109 | Su | Su_Hete |
| *Heliophobius argenteocinereus* | ZMB_Mam_107240 | Su | Su_Hete |
| *Chelemys macronyx* | NMW43360 | Su | Su_Noti |
| *Notoryctes typhlops* | NHMUK ZD 1981.1181 | Su | Su_Noto |
| *Notoryctes typhlops* | MNHN-ZM-MO 1893-473 | Su | Su_Noto |
| *Notoryctes typhlops* | ZMB_Mam_35694 | Su | Su_Noto |
| *Spalacopus cyanus* | ZMB_Mam_8317 | Su | Su_Octo |
| *Oryzorictes hova* | NHMUK ZD 1991.248 | Su | Su_Ory |
| *Oryzorictes hova* | MNHN-ZM-MO 1985-1624 | Su | Su_Ory |
| *Oryzorictes tetradactylus* | ZFMK MAM 1979.0155 | Su | Su_Ory |
| *Prometheomys schaposchnikowi* | MNHN_ZM_2012_24 | Su | Su_Prom |
| *Nannospalax ehrenbergi* | NMW65068 | Su | Su_Spal |
| *Rhizomys sumatrensis* | ZMB_Mam_21249 | Su | Su_Spal |
| *Spalax graecus* | NMW2158 | Su | Su_Spal |
| *Spalax microphthalmus* | ZMB_Mam_78545 | Su | Su_Spal |
| *Tachyoryctes splendens* | ZMB_Mam_72566 | Su | Su_Spal |
| *Euroscaptor mizura* | ZMB_Mam_103981 | Su | Su_Tal |
| *Mogera wogura* | ZMB_Mam_62455 | Su | Su_Tal |
| *Parascalops breweri* | NMW62569 | Su | Su_Tal |
| *Parascaptor leucura* | NHMUK ZE 1951.11.12.9 | Su | Su_Tal |
| *Scalopus aquaticus* | NHMUK ZE 1958.3.11.7 | Su | Su_Tal |
| *Talpa europaea* | ZMB_Mam_60682 | Su | Su_Tal |

| | | | |
|---|---|---|---|
| *Chlamyphorus truncatus* | ZMB_Mam_6007 | Su | Su_Xen |
| *Abrocoma budini* | NHMUK 1920.3.17.63 | Te | Else |
| *Abrocoma bennettii* | NMW23394 | Te | Else |
| *Caluromys philander* | ZMB_Mam_26760 | Te | Else |
| *Chinchilla lanigera* | ZMB_Mam_81126 | Te | Else |
| *Ctenodactylus gundi* | ZMB_Mam_71181 | Te | Else |
| *Dasyprocta punctata* | ZMB_Mam_72483 | Te | Else |
| *Dolichotis patagonum* | MNHN_AC_2000-827 | Te | Else |
| *Echinops telfairi* | ZMB_Mam_71612 | Te | Else |
| *Geogale aurita* | MNHN-ZM-MO1982-1001 | Te | Else |
| *Hystrix africaeaustralis* | ZMB_Mam_70881 | Te | Else |
| *Laonastes aenigmamus* | NHMUK ZD 1998.409 | Te | Else |
| *Lepus europaeus* | ZMB_Mam_70801 | Te | Else |
| *Microgale cowani* | ZMB_Mam_71614 | Te | Else |
| *Myrmecobius fasciatus* | ZMB_Mam_3121 | Te | Else |
| *Elephas maximus* | Confluences MHNL | Te | Else |
| *Ochotona princeps* | ZMB_Mam_93877 | Te | Else |
| *Ochotona rufescens* | MNHN_CG2000-409 | Te | Else |
| *Antechinus swainsonii* | IRSNB_21d | Te | Else |
| *Setifer setosus* | ZMB_Mam_44588 | Te | Else |
| *Sicista betulina* | NMW29531 | Te | Else |
| *Solenodon cubanus* | ZMB_Mam_2761 | Te | Else |
| *Sus scrofa* | ZMB_Mam_7975 | Te | Else |
| *Tayassu pecari* | ZMB_Mam_A.18.11 | Te | Else |
| *Tenrec ecaudatus* | NMW2432 | Te | Else |
| *Tupaia javanica* | ZMB_Mam_87169 | Te | Else |
| *Acinonyx jubatus* | MNHN_AC_1998.1981 | Te | Out_Ae_Chi |
| *Ailuropoda melanoleuca* | ZMB_Mam_17246 | Te | Out_Ae_Chi |
| *Canis lupus* | MNHN_CG_1996.2499 | Te | Out_Ae_Chi |
| *Crocuta crocuta* | ZMB_Mam_13295 | Te | Out_Ae_Chi |
| *Equus burchellii* | ZMB_Mam_15963 | Te | Out_Ae_Chi |
| *Helogale parvula* | ZMB_Mam_ 22986 | Te | Out_Ae_Chi |
| *Leptailurus serval* | ZMB_Mam_58145 | Te | Out_Ae_Chi |
| *Mustela eversmannii* | ZMB_Mam_12407 | Te | Out_Ae_Chi |
| *Mustela nivalis* | ZMB_Mam_ 56731 | Te | Out_Ae_Chi |
| *Diceros bicornis* | ZMB_Mam_32194 | Te | Out_Ae_Chi |
| *Tapirus indicus* | ZMB_Mam_4950 | Te | Out_Ae_Chi |
| *Civettictis civetta* | ZMB_Mam_68945 | Te | Out_Ae_Chi |
| *Vulpes vulpes* | ZMB_Mam_49955 | Te | Out_Ae_Chi |
| *Dicerorhinus sumatrensis* | ZMB_Mam_105847 | Te | Out_Ae_Chi |
| *Aotus azarae* | ZMB_Mam_35793 | Te | Out_Ae_Der |
| *Otolemur garnettii* | ZMB_Mam_5294 | Te | Out_Ae_Der |
| *Pan troglodytes* | MNHN_AC_1950.194 | Te | Out_Ae_Der |
| *Papio anubis* | ZMB_Mam_74869 | Te | Out_Ae_Der |
| *Aepyprymnus rufescens* | ZMB_Mam_35059 | Te | Out_Ae_Mars |
| *Bettongia gaimardi* | ZMB_Mam_60597 | Te | Out_Ae_Mars |
| *Bettongia lesueur* | NHMUK ZD 1858.10.16.3 | Te | Out_Ae_Mars |
| *Bettongia penicillata* | NHMUK ZD 1858.5.26.23 | Te | Out_Ae_Mars |
| *Hypsiprymnodon moschatus* | ZMB_Mam_78469 | Te | Out_Ae_Mars |
| *Potorous tridactylus* | MNHN-ZM-MO 1881-1148 | Te | Out_Ae_Mars |

| | | | |
|---|---|---|---|
| *Cephalophus silvicultor* | MNHN_AC_1981.1023 | Te | Out_Aq_Cet |
| *Dama dama* | ZMB_Mam_94752 | Te | Out_Aq_Cet |
| *Kobus ellipsiprymnus* | MNHN_AC_1887.1237 | Te | Out_Aq_Cet |
| *Lama glama* | NHMUK GERM 1860.e | Te | Out_Aq_Cet |
| *Moschus moschiferus* | ZMB_Mam_71049 | Te | Out_Aq_Cet |
| *Okapia johnstoni* | MNHN_AC_1978.27 | Te | Out_Aq_Cet |
| *Tragelaphus imberbis* | ZMB_Mam_48713 | Te | Out_Aq_Cet |
| *Tragulus napu* | ZMB_Mam_A 14 08 | Te | Out_Aq_Cet |
| *Vicugna vicugna* | ZMB_Mam_56204 | Te | Out_Aq_Cet |
| *Dendrohyrax arboreus* | ZMB_Mam_21098 | Te | Out_Aq_Sir_Su_Xen |
| *Dendrohyrax arboreus* | NHMUK 6.6.5.23 | Te | Out_Aq_Sir_Su_Xen |
| *Heterohyrax brucei* | ZMB_Mam_21479 | Te | Out_Aq_Sir_Su_Xen |
| *Procavia capensis* | ZMB_Mam_89416 | Te | Out_Aq_Sir_Su_Xen |
| *Calomyscus bailwardi* | NMW18256 | Te | Out_SciuAnoSpalProm |
| *Cricetomys gambianus* | ZMB_Mam_5371 | Te | Out_SciuAnoSpalProm |
| *Dicrostonyx groenlandicus* | ZMB_Mam_78542 | Te | Out_SciuAnoSpalProm |
| *Hyomys goliath* | ZMB_Mam_034358 | Te | Out_SciuAnoSpalProm |
| *Meriones meridianus* | ZMB_Mam_43005 | Te | Out_SciuAnoSpalProm |
| *Rhabdomys pumilio* | ZMB_Mam_86677 | Te | Out_SciuAnoSpalProm |
| *Tatera indica* | ZMB_Mam_72793 | Te | Out_SciuAnoSpalProm |
| *Uromys anak* | ZMB_Mam_34364 | Te | Out_SciuAnoSpalProm |
| *Petrodromus tetradactylus* | ZMB_Mam_84912 | Te | Out_Su_Chry_Ory |
| *Capromys pilorides* | ZMB_Mam_3202 | Te | Out_Su_Cten |
| *Thrichomys apereoides* | ZMB_Mam_8283 | Te | Out_Su_Cten |
| *Echimys chrysurus* | ZMB_Mam_8345 | Te | Out_Su_Cten |
| *Kannabateomys amblyonyx* | ZMB_Mam_7259 | Te | Out_Su_Cten |
| *Proechimys mincae* | ZMB_Mam_13543 | Te | Out_Su_Cten |
| *Lagurus lagurus* | NHMUK ZE 1962.3.21.2 | Te | Out_Su_Ello |
| *Chaetodipus formosus* | MNHN-ZM-MO 1914-35B | Te | Out_Su_Geo |
| *Heteromys desmarestianus* | ZFMK MAM 1981.1464 | Te | Out_Su_Geo |
| *Perognathus longimembris* | NMW61838 | Te | Out_Su_Geo |
| *Perognathus fasciatus* | ZMB_Mam_5071 | Te | Out_Su_Geo |
| *Petromus typicus* | ZMB_Mam_105768 | Te | Out_Su_Hete |
| *Thryonomys swinderianus* | ZMB_Mam_72392 | Te | Out_Su_Hete |
| *Thryonomys gregorianus* | NHMUK 76.193 | Te | Out_Su_Hete |
| *Abrothrix olivaceus* | MNHN-ZM-MO_1884-1293 | Te | Out_Su_Noti |
| *Abrothrix longipilis* | NHMUK ZD 1858.9.6.5 | Te | Out_Su_Noti |
| *Echymipera clara* | ZMB_Mam_15764 | Te | Out_Su_Noto |
| *Perameles nasuta* | IRSNB_39 | Te | Out_Su_Noto |
| *Octodon degus* | ZMB_Mam_8321 | Te | Out_Su_Octo |
| *Atelerix albiventris* | ZMB_Mam_5810 | Te | Out_Su_Tal |
| *Echinosorex gymnura* | ZMB_Mam_72232 | Te | Out_Su_Tal |
| *Hemiechinus auritus* | NMW41180 | Te | Out_Su_Tal |
| *Hylomys megalotis* | NHMUK ZD 1999.47 | Te | Out_Su_Tal |
| *Hylomys suillus* | NHMUK ZE 1960.8.4.7 | Te | Out_Su_Tal |
| *Paraechinus hypomelas* | NMW15184 | Te | Out_Su_Tal |
| *Uropsilus soricipes* | NMW64409 | Te | Out_Su_Tal |

**Footnotes.** Lifestyles: Ae, aerial; Aq, fully aquatic; Su, subterranean, Te, Terrestrial. Clades: each specialised clade is referred to with the abbreviation of the corresponding lifestyle followed by a unique abbreviation; the terrestrial most closely related species of each specialised clade is referred to as “Out_*the abbreviation of the specialised clade* (note that some of these correspond to several specialised clades); Else, other, more distantly related terrestrial families. Museum collections: IRSNB, Institut royal des Sciences naturelles de Bruxelles; MHNL, Musée des Confluences, Lyon, France; MNHN, Muséum national d’Histoire naturelle, Paris, France; NHMUK, Natural History Museum, London, UK; NMW, Naturhistorisches Museum Wien, Austria; ZFMK MAM, Zoologische Forschungsmuseum Alexander Koenig, Bonn, Germany; ZMB_Mam, Mammals collection of the Museum für Naturkunde, Berlin.

### Supplementary Table S2. Descriptive statistics

Descriptive statistics

| | | Mean | Min | Max |
|---|---|---|---|---|
| Vertebral mean Cg | | | | |
| | Terrestrial | 35.220 | 17.789 | 62.674 |
| | Aerial | 33.164 | 14.921 | 45.912 |
| | Aquatic | 62.425 | 58.249 | 71.983 |
| | Subterranean | 30.556 | 17.289 | 51.209 |
| Connectivity | | | | |
| | Terrestrial | 4958.585 | 0.000 | 298141.750 |
| | Aerial | 19.134 | 0.000 | 98.125 |
| | Aquatic | 100620.699 | 3500.000 | 711702.375 |
| | Subterranean | 82.298 | 2.500 | 1191.625 |
| BV.TV | | | | |
| | Terrestrial | 0.228 | 0.000 | 0.533 |
| | Aerial | 0.212 | 0.000 | 0.425 |
| | Aquatic | 0.423 | 0.309 | 0.703 |
| | Subterranean | 0.203 | 0.029 | 0.489 |
| Humeral mean Cg | | | | |
| | Terrestrial | 60.664 | 37.513 | 82.480 |
| | Aerial | 53.130 | 30.213 | 62.519 |
| | Aquatic | 78.030 | 49.205 | 99.619 |
| | Subterranean | 61.914 | 42.445 | 80.032 |
| CSS | | | | |
| | Terrestrial | 2.070 | 1.121 | 5.252 |
| | Aerial | 1.268 | 1.031 | 1.660 |
| | Aquatic | 1.953 | 1.281 | 2.873 |
| | Subterranean | 3.157 | 1.242 | 10.837 |
| DE | | | | |
| | Terrestrial | 4.819 | 3.065 | 7.026 |
| | Aerial | 7.200 | 5.336 | 9.924 |
| | Aquatic | 2.386 | 0.847 | 3.681 |
| | Subterranean | 3.364 | 1.162 | 5.715 |

### Supplementary Results S1. AN(C)OVAs

AN(C)OVAs detailed outputs.

Lifestyle abbreviations: Ae, Aerial; Aq, Aquatic; Te, Terrestrial; Su, Subterranean.

### 1. Vertebral parameters

1.1. Whole dataset

**Vertebral mean Cg ~ log(Vertebral_centrum_length) + Lifestyle**

| Pagel's lambda |
|---|
| 0.5515288 |

Coefficients:

| | Value | Std.Error | t-value | p-value |
|---|---|---|---|---|
| (Intercept) | 15.967524 | 3.752081 | 4.255645 | 0.0000 |
| log(VertL) | 9.362762 | 0.825957 | 11.335656 | **0.0000** |

Multiple Comparisons of Means: Tukey Contrasts

| | Value | Std.Error | z-value | Pr(>\|z\|) |
|---|---|---|---|---|
| Aq - Ae | 10.6641 | 4.0266 | 2.648 | **0.03541** |
| Su - Ae | 0.4067 | 2.3187 | -0.175 | 0.99786 |
| Te - Ae | -1.8826 | 2.0900 | -0.901 | 0.78737 |
| Su - Aq | -11.0708 | 3.7476 | -2.954 | **0.01438** |
| Te - Aq | -12.5466 | 3.4651 | -3.621 | **0.00141** |
| Te - Su | -1.4758 | 1.4645 | -1.008 | 0.72439 |

Vertebral mean Cg ~ log(Vertrebral_centrum_length)

| | Value | Std.Error | t-value | p-value |
|---|---|---|---|---|
| (Intercept) | 14.10409 | 4.159763 | 3.390599 | 9e-04 |
| log(VertL) | 10.02336 | 0.823731 | 12.168250 | 0.0000 |

Nagelkerke pseudo $R^2$: 0.5869862

---

**log(Connectivity) ~ log(Vertebral_centrum_length) + Lifestyle**

| Pagel's lambda |
|---|
| 0.55818 |

Coefficients:

| | Value | Std.Error | t-value | p-value |
|---|---|---|---|---|
| (Intercept) | -0.760339 | 0.7991580 | -0.951425 | 0.3427 |
| log(VertL) | 1.771254 | 0.1602438 | 11.053496 | **0.0000** |

Multiple Comparisons of Means: Tukey Contrasts

| | Value | Std.Error | t-value | p-value |
|---|---|---|---|---|
| Aq - Ae | 3.6395 | 0.7825 | 4.651 | **< 0.001** |
| Su - Ae | 1.5695 | 0.4958 | 3.165 | **0.00736** |
| Te - Ae | 0.6036 | 0.4470 | 1.350 | 0.50519 |
| Su - Aq | -2.0701 | 0.7179 | -2.883 | 0.01831 |
| Te - Aq | -3.0360 | 0.6452 | -4.706 | **< 0.001** |
| Te - Su | -0.9659 | 0.3133 | -3.083 | **0.00969** |

log(Connectivity) ~ log(Vertebral_centrum_length)

| | Value | Std.Error | t-value | p-value |
|---|---|---|---|---|
| (Intercept) | -0.1872968 | 1.0885475 - | 0.172061 | 0.8636 |
| log(VertL) | 1.8862857 | 0.1711185 | 11.023273 | 0.0000 |

Nagelkerke pseudo $R^2$: 0.4824922

---

**BV.TV ~ log(Vertrebral_centrum_length) + Lifestyle**

| Pagel's lambda |
|---|
| 0.3461863 |

Coefficients

| | Value | Std.Error | t-value | p-value |
|---|---|---|---|---|
| (Intercept) | 0.08123194 | 0.03981464 | 2.040253 | 0.0428 |
| log(VertL) | 0.07407308 | 0.00958539 | 7.727704 | **0.0000** |

Multiple Comparisons of Means: Tukey Contrasts

| | Value | Std.Error | t-value | p-value |
|---|---|---|---|---|
| Aq - Ae | 0.0233287 | 0.0422055 | 0.553 | 0.941 |
| Su - Ae | -0.0009192 | 0.0272091 | -0.034 | 1.000 |
| Te - Ae | -0.0337508 | 0.0244241 | -1.382 | 0.487 |
| Su - Aq | -0.0242480 | 0.0396518 | -0.612 | 0.922 |
| Te - Aq | -0.0570795 | 0.0347457 | -1.643 | 0.333 |
| Te - Su | -0.0328316 | 0.0188409 | -1.743 | 0.281 |

BV.TV ~ log(Vertebral_centrum_length)

| | Value | Std.Error | t-value | p-value |
|---|---|---|---|---|
| (Intercept) | 0.06783887 | 0.04113441 | 1.649200 | 0.1009 |
| log(VertL) | 0.07307459 | 0.00891487 | 8.196933 | 0.0000 |

Nagelkerke pseudo $R^2$: 0.3778663

### 1.2. Terrestrial pruned to match Aquatic size

**Vertebral Mean Cg ~ log(Vertebral_centrum_length) + Lifestyle**

| Pagel's lambda |
|---|

| |
|---|
| 0.6178141 |

Coefficients

| | Value | Std.Error | t-value | p-value |
|---|---|---|---|---|
| (Intercept) | 10.456846 | 5.730015 | 1.824925 | 0.0716 |
| log(VertL) | 10.607906 | 1.268990 | 8.359330 | **0.0000** |
| Te-Aq | 13.363764 | 3.322355 | 4.022377 | **0.0001** |

**log(Connectivity) ~ log(Vertebral_centrum_length) + Lifestyle**

| Pagel's lambda |
|---|
| 0.5547799 |

Coefficients

| | Value | Std.Error | t-value | p-value |
|---|---|---|---|---|
| (Intercept) | 1.7962133 | 1.2443517 | 1.443493 | 0.1518 |
| log(VertL) | 1.4809046 | 0.2659399 | 5.568568 | **0.0000** |
| Te-Aq | 2.8420086 | 0.6676666 | 4.256629 | **0.0000** |

#### 1.3. Terrestrial pruned to match Aerial and Subterranean Size

**log(Connectivity) ~ log(Vertebral_centrum_length) + Lifestyle**

| Pagel's lambda |
|---|
| 0.4313344 |

Coefficients

| | Value | Std.Error | t-value | p-value |
|---|---|---|---|---|
| (Intercept) | 0.731928 | 0.6578026 | 1.112687 | 0.2679 |
| log(VertL) | 1.252803 | 0.2184246 | 5.735632 | **0.0000** |
| Te-Ae | -0.525197 | 0.4018929 | -1.306808 | 0.1936 |
| Te-Su | 0.995753 | 0.2857582 | 3.484600 | **0.0007** |

### 2. Humeral parameters

#### 2.1. Whole dataset

**Humeral mean Cg ~ log(mean_total_cross-sectional_area) + Lifestyle**

| Pagel's lambda |
|---|
| 0.64127 |

Coefficients

| | Value | Std.Error | t-value | p-value |
|---|---|---|---|---|
| (Intercept) | 52.27987 | 5.378621 | 9.719939 | 0.0000 |
| log(MeanArea) | 1.32066 | 0.553781 | 2.384808 | **0.0184** |

Multiple Comparisons of Means: Tukey Contrasts

| | Value | Std.Error | t-value | p-value |
|---|---|---|---|---|
| Aq - Ae | 15.7933 | 6.2245 | 2.537 | **0.047** |
| Su - Ae | 5.9057 | 3.4534 | 1.710 | 0.291 |
| Te - Ae | 5.0618 | 3.0864 | 1.640 | 0.328 |
| Su - Aq | -9.8876 | 5.6649 | -1.745 | 0.274 |
| Te - Aq | -10.7315 | 5.4111 | -1.983 | 0.174 |
| Te - Su | -0.8439 | 2.0089 | -0.420 | 0.972 |

Humeral mean Cg ~ log(mean_total_cross-sectional_area)

| | Value | Std.Error | t-value | p-value |
|---|---|---|---|---|
| (Intercept) | 56.06108 | 5.422449 | 10.338702 | 0.000 |
| log(MeanArea) | 1.79860 | 0.537809 | 3.344315 | 0.001 |

Nagelkerke pseudo $R^2$: 0.09309527

---

**log(CSS) ~ log(mean_total_cross-sectional_area) + Lifestyle**

| Pagel's lambda |
|---|
| 0.6544086 |

Coefficients

| | Value | Std.Error | t-value | p-value |
|---|---|---|---|---|
| (Intercept) | 0.2291447 | 0.22828617 | 1.003761 | 0.3171 |
| log(MeanArea) | 0.0302043 | 0.02320436 | 1.301667 | <u>0.1951</u> |

Multiple Comparisons of Means: Tukey Contrasts

| | Value | Std.Error | t-value | p-value |
|---|---|---|---|---|
| Aq - Ae | 0.39033 | 0.26194 | 1.490 | 0.41350 |
| Su - Ae | 0.69558 | 0.14557 | 4.778 | **< 0.001** |
| Te - Ae | 0.46138 | 0.13016 | 3.545 | **0.00174** |
| Su - Aq | 0.30525 | 0.23818 | 1.282 | 0.54551 |
| Te - Aq | 0.07106 | 0.22756 | 0.312 | 0.98811 |
| Te - Su | -0.23419 | 0.08422 | -2.781 | **0.02354** |

log(CSS) ~ log(mean_total_cross-sectional_area)

| | Value | Std.Error | t-value | p-value |
|---|---|---|---|---|
| (Intercept) | 0.6431275 | 0.29040728 | 2.214571 | 0.0283 |
| log(MeanArea) | 0.0421687 | 0.02446698 | 1.723495 | <u>0.0869</u> |

Nagelkerke pseudo $R^2$: 0.02652733

---

**log(DE) ~ log(mean_total_cross-sectional_area) + Lifestyle**

| Pagel's lambda |
| --- |
| 0.9732087 |

Coefficients

| | Value | Std.Error | t-value | p-value |
| --- | --- | --- | --- | --- |
| (Intercept) | 2.0175333 | 0.18358951 | 10.989371 | 0.0000 |
| log(MeanArea) | -0.0419341 | 0.01234671 - | 3.396376 | **9e-04** |

Multiple Comparisons of Means: Tukey Contrasts

| | Value | Std.Error | t-value | p-value |
| --- | --- | --- | --- | --- |
| Aq - Ae | -1.15655 | 0.17342 | -6.669 | **< 0.001** |
| Su - Ae | -0.68674 | 0.09803 | -7.006 | **< 0.001** |
| Te - Ae | -0.42660 | 0.09062 | -4.708 | **< 0.001** |
| Su - Aq | 0.46981 | 0.15338 | 3.063 | **0.00976** |
| Te - Aq | 0.72995 | 0.14886 | 4.904 | **< 0.001** |
| Te - Su | 0.26014 | 0.04426 | 5.878 | **< 0.001** |

log(DE) ~ log(mean_total_cross-sectional_area)

| | Value | Std.Error | t-value | p-value |
| --- | --- | --- | --- | --- |
| (Intercept) | 1.6223822 | 0.2285688 | 7.098002 | 0.0000 |
| log(MeanArea) | -0.0627009 | 0.0144669 | -4.334094 | 0.0000 |

Nagelkerke pseudo $R^2$: 0.085116

### 2.2. Terrestrial pruned to match Aerial and Subterranean Size

**log(CSS) ~ log(mean_total_cross-sectional_area) + Lifestyle**

| Pagel's lambda |
| --- |
| 0.5932886 |

Coefficients

| | Value | Std.Error | t-value | p-value |
| --- | --- | --- | --- | --- |
| (Intercept) | 0.6612625 | 0.20331205 | 3.252451 | 0.0015 |
| log(MeanArea) | 0.0640891 | 0.03335165 | 1.921618 | 0.0569 |
| Te-Ae | -0.5146113 | 0.12943284 - | 3.975894 | **0.0001** |
| Te-Su | 0.2052093 | 0.08697299 | 2.359460 | **0.0198** |

**log(DE) ~ log(mean_total_cross-sectional_area) + Lifestyle**

| Pagel's lambda |
| --- |
| 0.9837882 |

Coefficients

| | Value | Std.Error | t-value | p-value |
| --- | --- | --- | --- | --- |

| | | | | |
|---|---|---|---|---|
| (Intercept) | 1.5611948 | 0.17930662 | 8.706844 | 0.000 |
| log(MeanArea) | -0.0277517 | 0.01609240 - | 1.724522 | 0.087 |
| Te-Ae | 0.4323139 | 0.09628532 | 4.489925 | **0.000** |
| Te-Su | -0.2558572 | 0.04572239 | -5.595885 | **0.000** |

**Supplementary Results S2. Phenograms.**

Phenograms depicting the reconstructed evolution of each trait among specialised clades. For clade name abbreviations, see Supplementary Table S1. Time in millions of years. Trait are either size-corrected and/or without units. Function phenogram (phytools package (Revell 2012)).

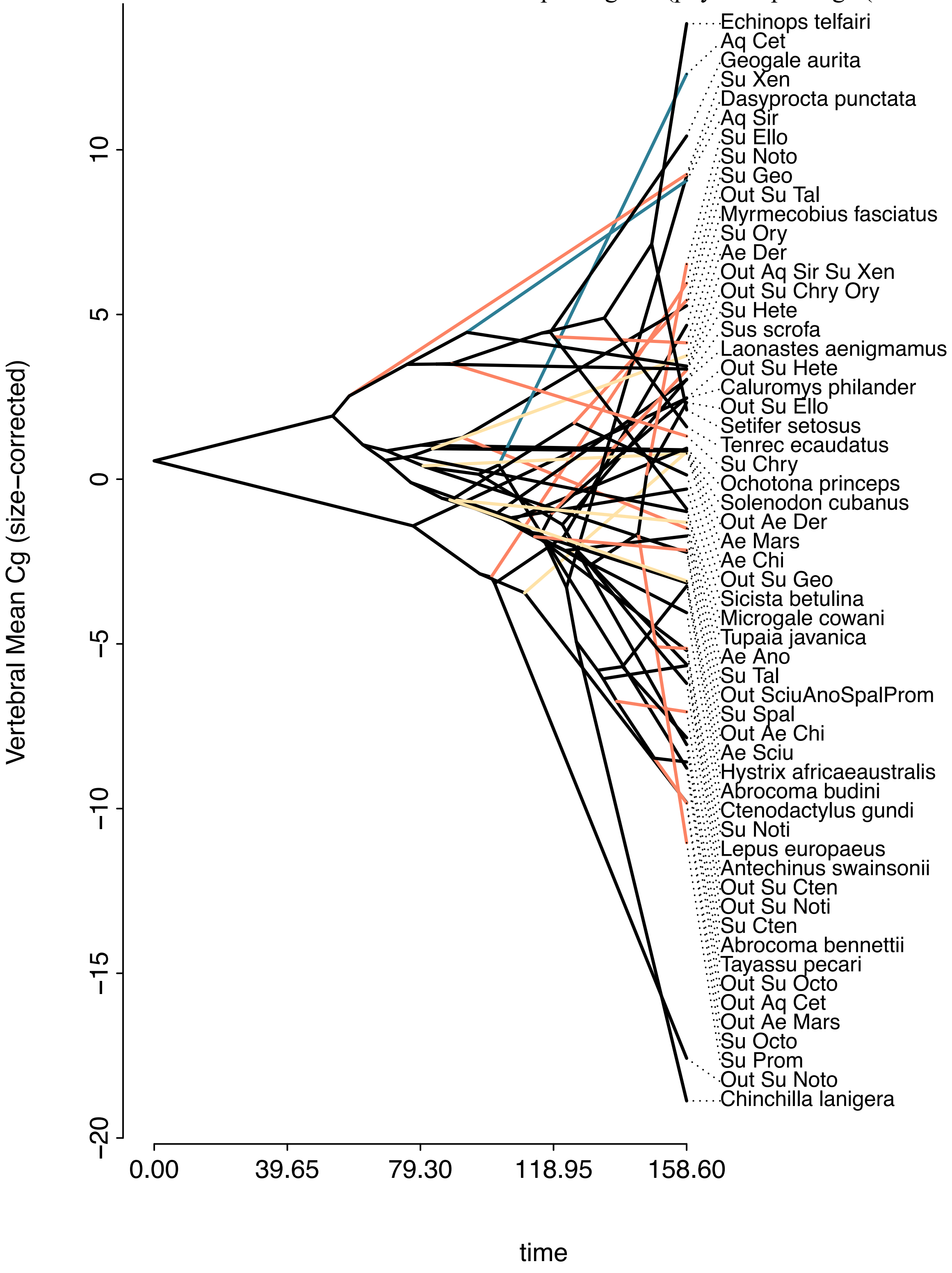

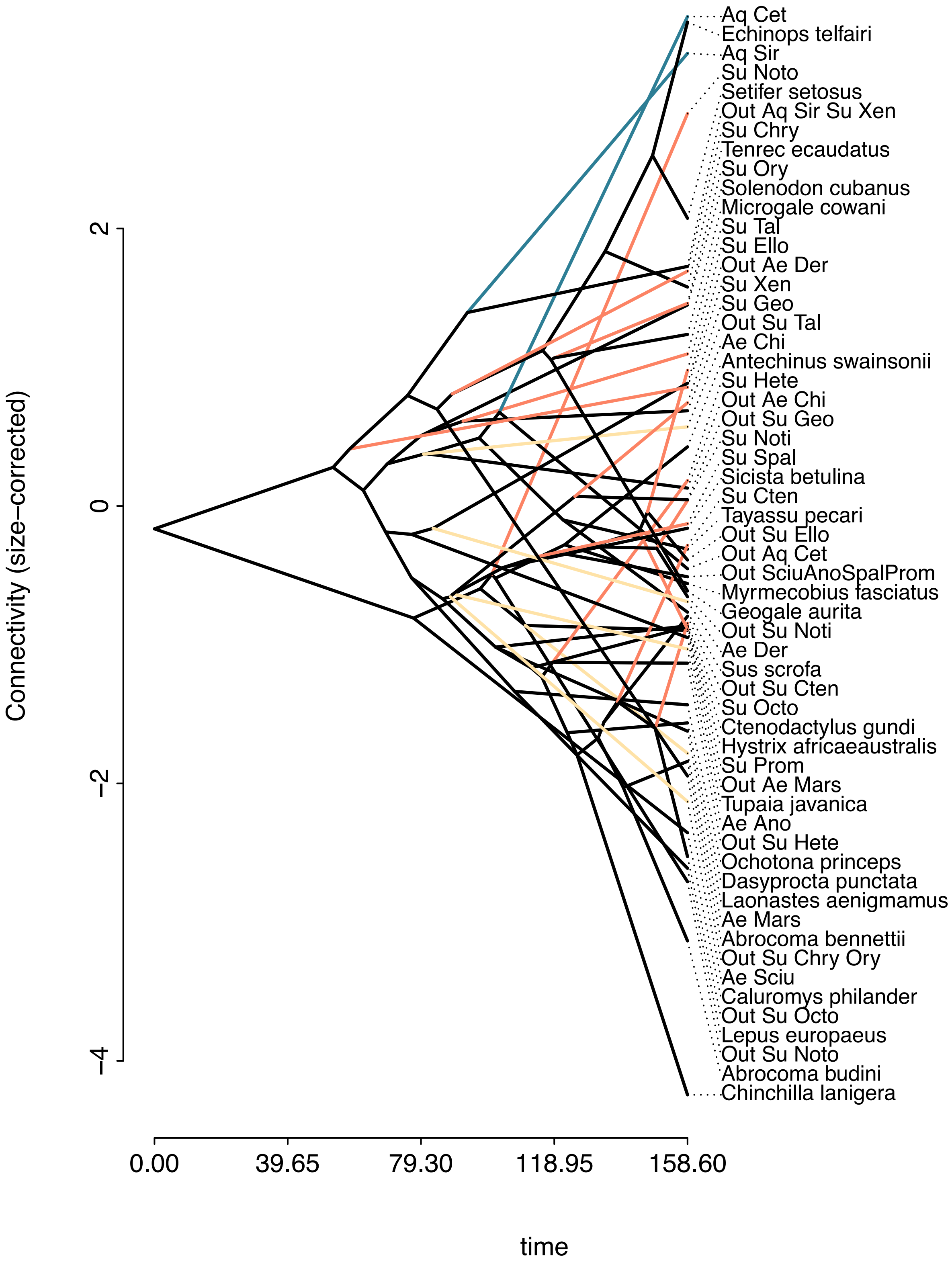
Connectivity (size−corrected)
2
0
−2
−4
0.00
39.65
79.30
118.95
158.60
time
Aq Cet
Echinops telfairi
Aq Sir
Su Noto
Setifer setosus
Out Aq Sir Su Xen
Su Chry
Tenrec ecaudatus
Su Ory
Solenodon cubanus
Microgale cowani
Su Tal
Su Ello
Out Ae Der
Su Xen
Su Geo
Out Su Tal
Ae Chi
Antechinus swainsonii
Su Hete
Out Ae Chi
Out Su Geo
Su Noti
Su Spal
Sicista betulina
Su Cten
Tayassu pecari
Out Su Ello
Out Aq Cet
Out SciuAnoSpalProm
Myrmecobius fasciatus
Geogale aurita
Out Su Noti
Ae Der
Sus scrofa
Out Su Cten
Su Octo
Ctenodactylus gundi
Hystrix africaeaustralis
Su Prom
Out Ae Mars
Tupaia javanica
Ae Ano
Out Su Hete
Ochotona princeps
Dasyprocta punctata
Laonastes aenigmamus
Ae Mars
Abrocoma bennettii
Out Su Chry Ory
Ae Sciu
Caluromys philander
Out Su Octo
Lepus europaeus
Out Su Noto
Abrocoma budini
Chinchilla lanigera

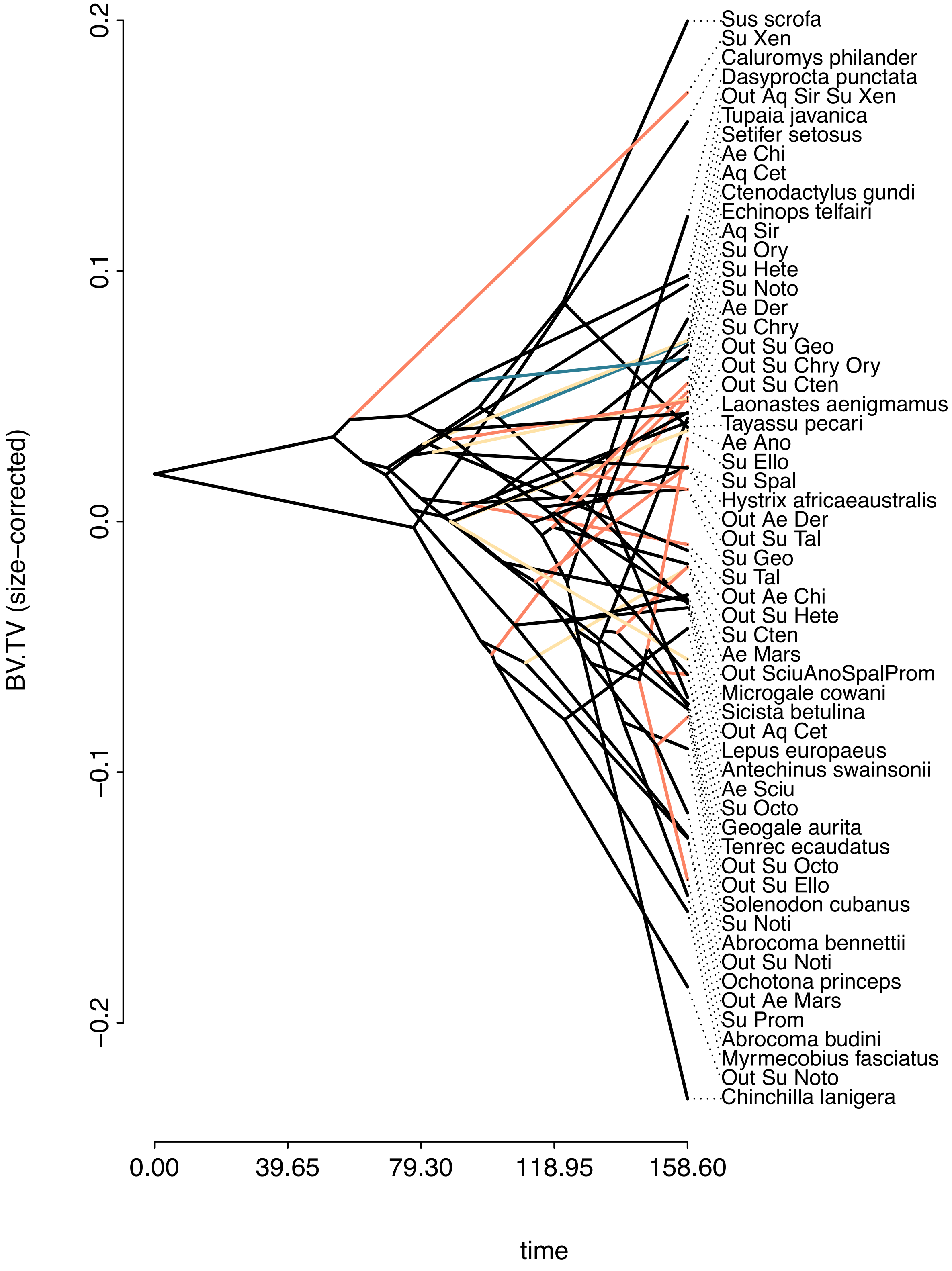
BV.TV (size−corrected)
0.2
0.1
0.0
−0.1
−0.2
0.00
39.65
79.30
118.95
158.60
time
Sus scrofa
Su Xen
Caluromys philander
Dasyprocta punctata
Out Aq Sir Su Xen
Tupaia javanica
Setifer setosus
Ae Chi
Aq Cet
Ctenodactylus gundi
Echinops telfairi
Aq Sir
Su Ory
Su Hete
Su Noto
Ae Der
Su Chry
Out Su Geo
Out Su Chry Ory
Out Su Cten
Laonastes aenigmamus
Tayassu pecari
Ae Ano
Su Ello
Su Spal
Hystrix africaeaustralis
Out Ae Der
Out Su Tal
Su Geo
Su Tal
Out Ae Chi
Out Su Hete
Su Cten
Ae Mars
Out SciuAnoSpalProm
Microgale cowani
Sicista betulina
Out Aq Cet
Lepus europaeus
Antechinus swainsonii
Ae Sciu
Su Octo
Geogale aurita
Tenrec ecaudatus
Out Su Octo
Out Su Ello
Solenodon cubanus
Su Noti
Abrocoma bennettii
Out Su Noti
Ochotona princeps
Out Ae Mars
Su Prom
Abrocoma budini
Myrmecobius fasciatus
Out Su Noto
Chinchilla lanigera

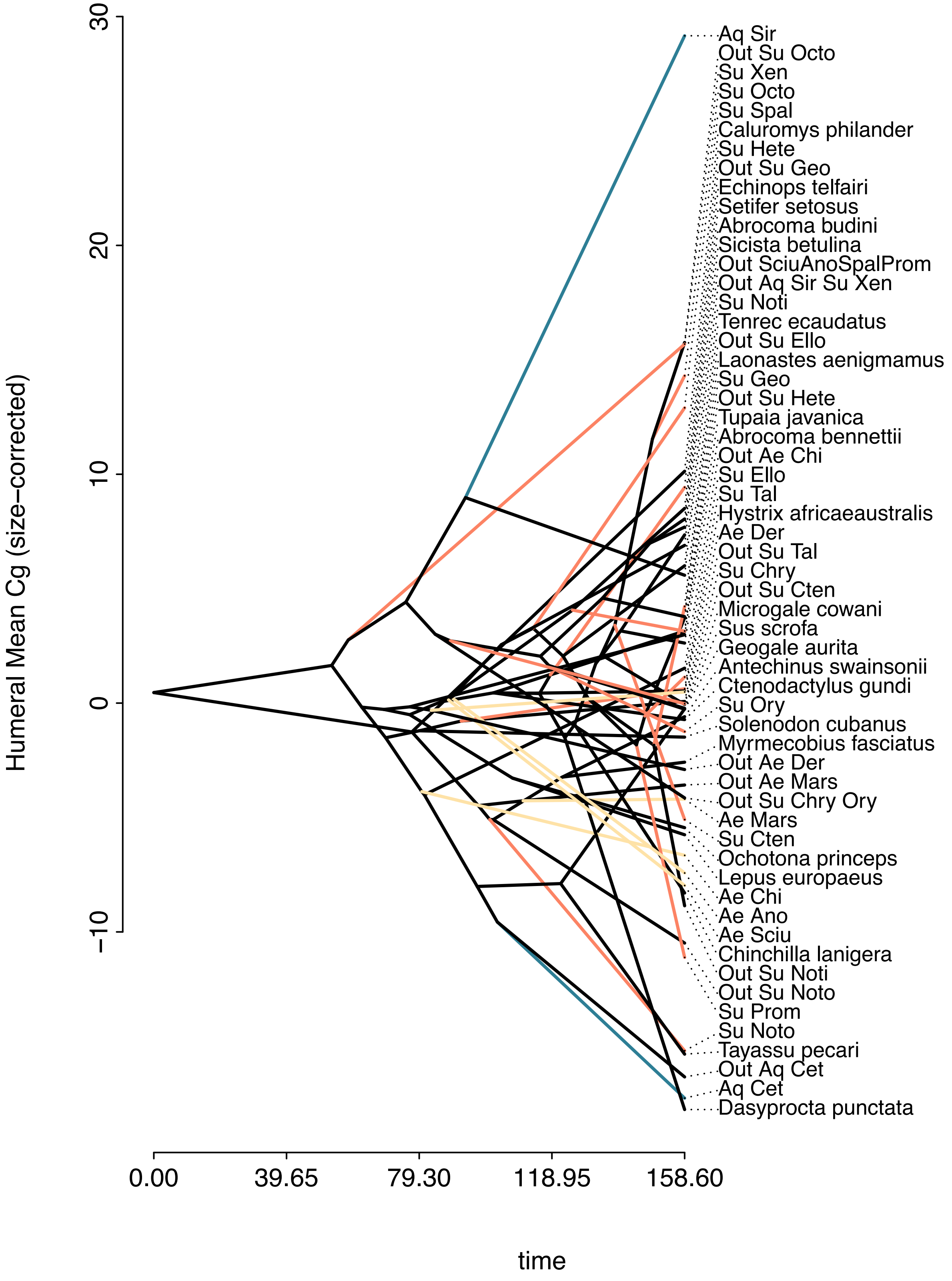
Humeral Mean Cg (size-corrected)
30
20
10
0
−10
time
0.00
39.65
79.30
118.95
158.60
Aq Sir
Out Su Octo
Su Xen
Su Octo
Su Spal
Caluromys philander
Su Hete
Out Su Geo
Echinops telfairi
Setifer setosus
Abrocoma budini
Sicista betulina
Out SciuAnoSpalProm
Out Aq Sir Su Xen
Su Noti
Tenrec ecaudatus
Out Su Ello
Laonastes aenigmamus
Su Geo
Out Su Hete
Tupaia javanica
Abrocoma bennettii
Out Ae Chi
Su Ello
Su Tal
Hystrix africaeaustralis
Ae Der
Out Su Tal
Su Chry
Out Su Cten
Microgale cowani
Sus scrofa
Geogale aurita
Antechinus swainsonii
Ctenodactylus gundi
Su Ory
Solenodon cubanus
Myrmecobius fasciatus
Out Ae Der
Out Ae Mars
Out Su Chry Ory
Ae Mars
Su Cten
Ochotona princeps
Lepus europaeus
Ae Chi
Ae Ano
Ae Sciu
Chinchilla lanigera
Out Su Noti
Out Su Noto
Su Prom
Su Noto
Tayassu pecari
Out Aq Cet
Aq Cet
Dasyprocta punctata

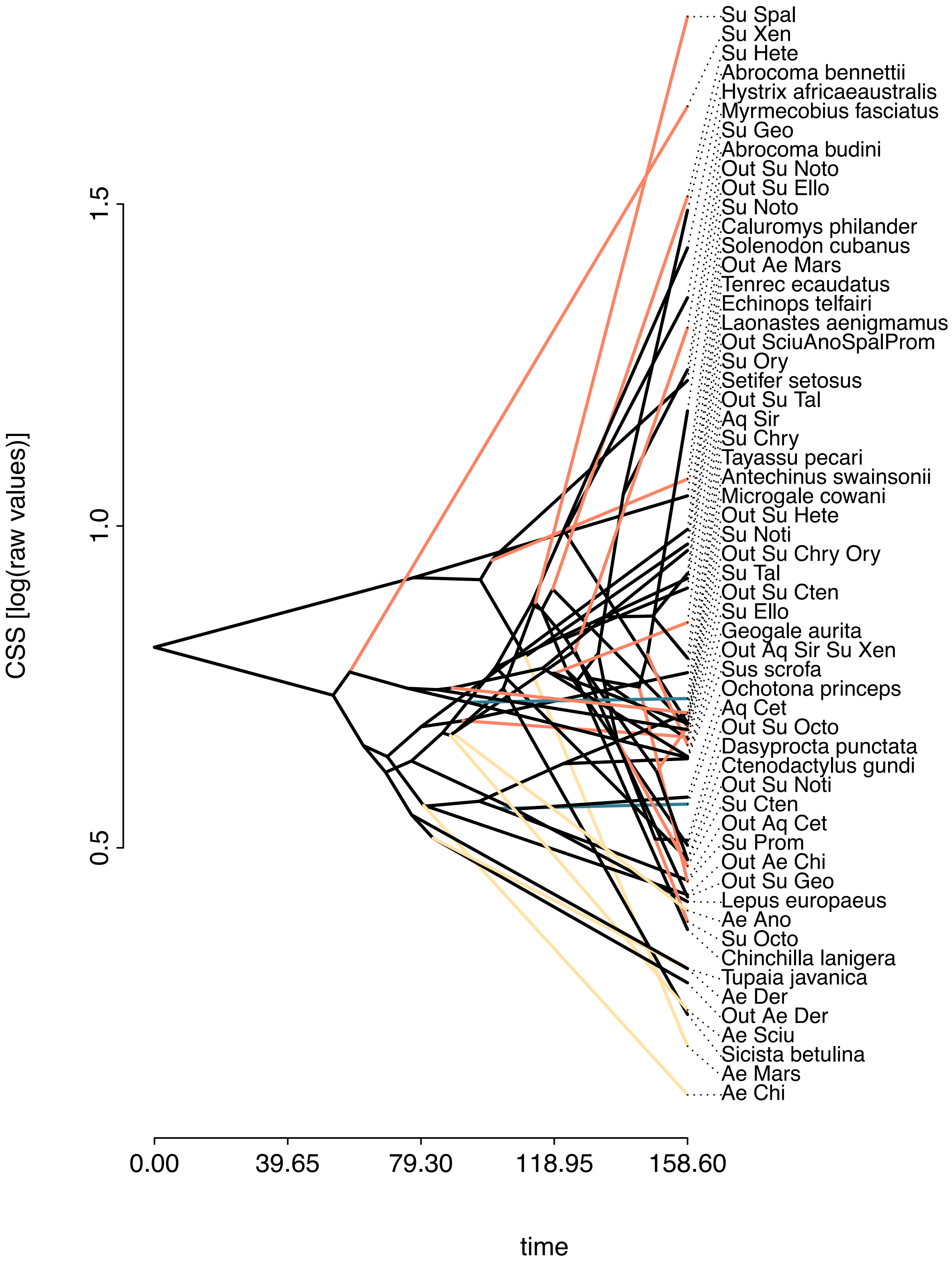
Su Spal
Su Xen
Su Hete
Abrocoma bennettii
Hystrix africaeaustralis
Myrmecobius fasciatus
Su Geo
Abrocoma budini
Out Su Noto
Out Su Ello
Su Noto
Caluromys philander
Solenodon cubanus
Out Ae Mars
Tenrec ecaudatus
Echinops telfairi
Laonastes aenigmamus
Out SciuAnoSpalProm
Su Ory
Setifer setosus
Out Su Tal
Aq Sir
Su Chry
Tayassu pecari
Antechinus swainsonii
Microgale cowani
Out Su Hete
Su Noti
Out Su Chry Ory
Su Tal
Out Su Cten
Su Ello
Geogale aurita
Out Aq Sir Su Xen
Sus scrofa
Ochotona princeps
Aq Cet
Out Su Octo
Dasyprocta punctata
Ctenodactylus gundi
Out Su Noti
Su Cten
Out Aq Cet
Su Prom
Out Ae Chi
Out Su Geo
Lepus europaeus
Ae Ano
Su Octo
Chinchilla lanigera
Tupaia javanica
Ae Der
Out Ae Der
Ae Sciu
Sicista betulina
Ae Mars
Ae Chi
CSS [log(raw values)]
1.5
1.0
0.5
0.00
39.65
79.30
118.95
158.60
time

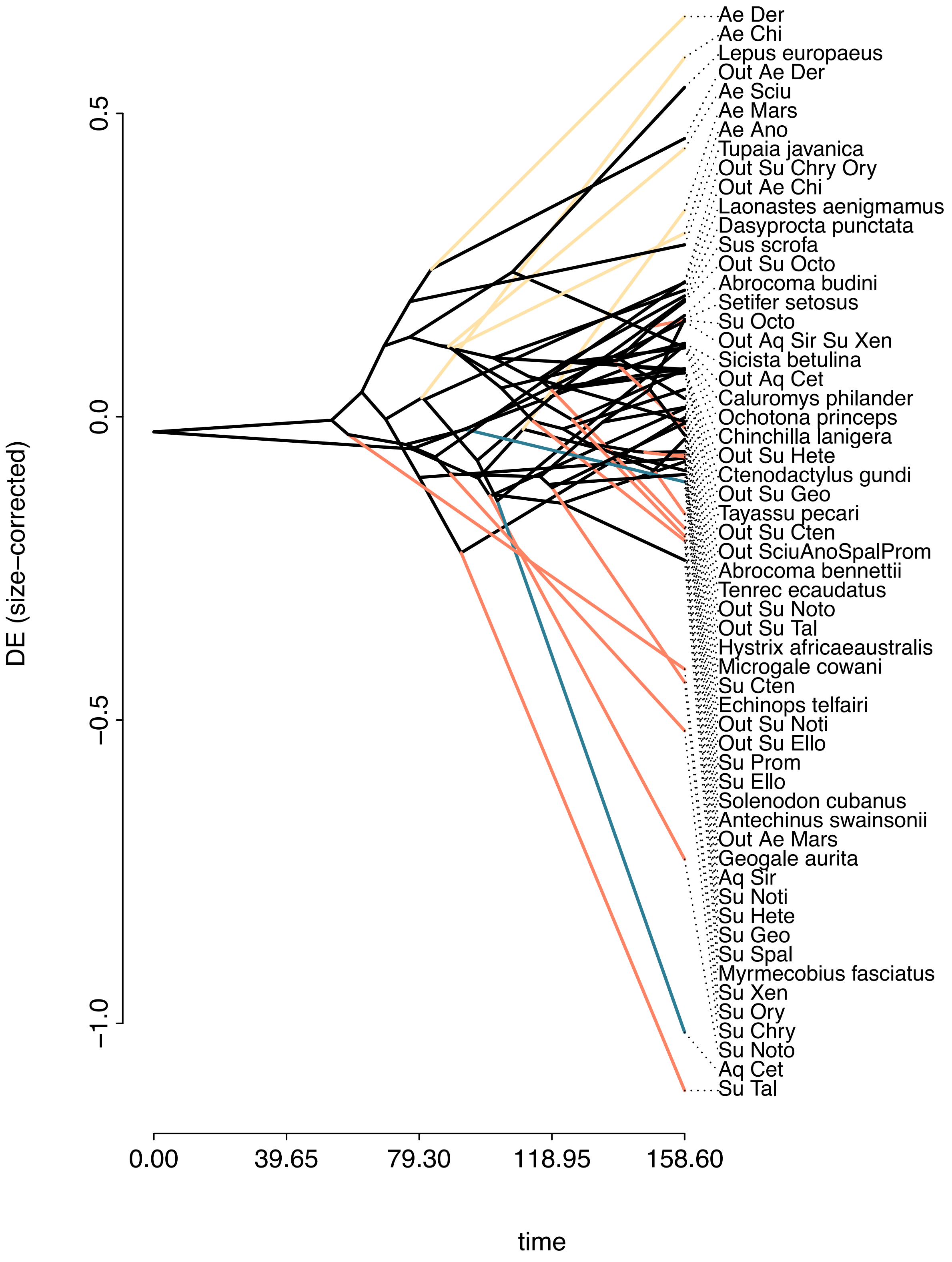

DE (size-corrected)
0.5
0.0
−0.5
−1.0
0.00
39.65
79.30
118.95
158.60
time
Ae Der
Ae Chi
Lepus europaeus
Out Ae Der
Ae Sciu
Ae Mars
Ae Ano
Tupaia javanica
Out Su Chry Ory
Out Ae Chi
Laonastes aenigmamus
Dasyprocta punctata
Sus scrofa
Out Su Octo
Abrocoma budini
Setifer setosus
Su Octo
Out Aq Sir Su Xen
Sicista betulina
Out Aq Cet
Caluromys philander
Ochotona princeps
Chinchilla lanigera
Out Su Hete
Ctenodactylus gundi
Out Su Geo
Tayassu pecari
Out Su Cten
Out SciuAnoSpalProm
Abrocoma bennettii
Tenrec ecaudatus
Out Su Noto
Out Su Tal
Hystrix africaeaustralis
Microgale cowani
Su Cten
Echinops telfairi
Out Su Noti
Out Su Ello
Su Prom
Su Ello
Solenodon cubanus
Antechinus swainsonii
Out Ae Mars
Geogale aurita
Aq Sir
Su Noti
Su Hete
Su Geo
Su Spal
Myrmecobius fasciatus
Su Xen
Su Ory
Su Chry
Su Noto
Aq Cet
Su Tal

### Supplementary Results S3. Phylogenetic signal.

Phylogenetic signal among terrestrial species.

Sample size range from 82 to 92 (depending on the trait), for all species analysed individually and is 37 when terrestrial sister-groups (TSG) are aggregated. Mapping (function contmap, phytools package (Revell 2012) for the Connectivity and diaphysis elongation (DE), the only traits for which a significant signal was found when the TSG are aggregated, is also displayed.

**Vertebral mean Cg (size-corrected)**

All species individually

Phylogenetic signal lambda: 0.46183
logL(lambda): -277.557
LR(lambda=0): 10.6376
P-value (based on LR test): **0.00110812**

TSG aggregated

Phylogenetic signal lambda: 0.204127
logL(lambda): -122.523
LR(lambda=0): 0.522751
P-value (based on LR test): 0.46967

**BV.TV (size-corrected)**

All species individually

Phylogenetic signal lambda: 0.258965
logL(lambda): 88.6745
LR(lambda=0): 5.24641
P-value (based on LR test): **0.021992**

TSG aggregated

Phylogenetic signal lambda: 7.07461e-05
logL(lambda): 34.6072
LR(lambda=0): -0.000517764
P-value (based on LR test): 1

**Connectivity (size-corrected)**

All species individually

Phylogenetic signal lambda: 0.597226
logL(lambda): -170.849
LR(lambda=0): 13.4913
P-value (based on LR test): **0.000239667**

TSG aggregated

Phylogenetic signal lambda: 0.844203
logL(lambda): -62.8934
LR(lambda=0): 11.912
P-value (based on LR test): **0.000557721**

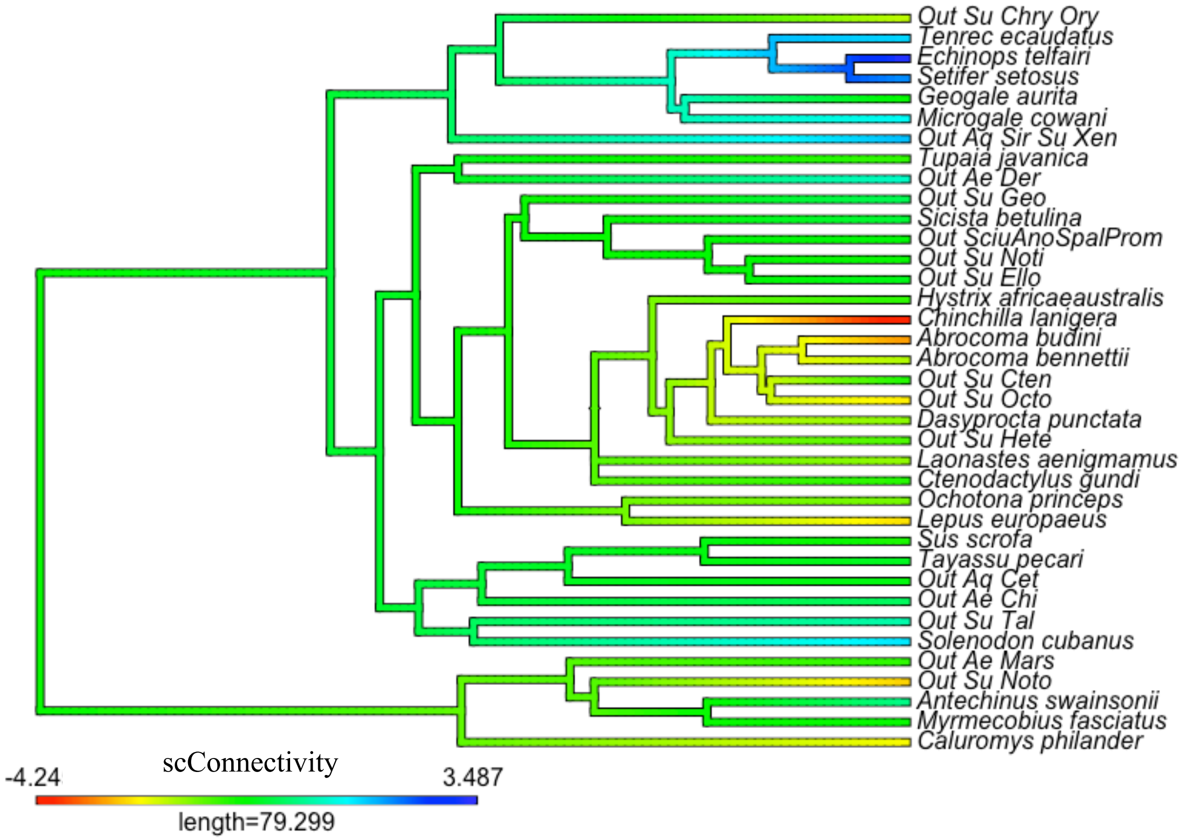


**Humeral mean Cg (size-corrected)**

All species individually

Phylogenetic signal lambda: 0.551538
logL(lambda): -295.984
LR(lambda=0): 6.7443
P-value (based on LR test): **0.00940477**

TSG aggregated

Phylogenetic signal lambda: 7.07461e-05
logL(lambda): -126.008
LR(lambda=0): -0.000977227
P-value (based on LR test): 1

**log(CSS)**

All species individually

Phylogenetic signal lambda: 0.465252
logL(lambda): -19.5601
LR(lambda=0): 11.6607
P-value (based on LR test): **0.000638344**

TSG aggregated

Phylogenetic signal lambda: 0.114551
logL(lambda): -10.927

LR(lambda=0): 0.123667
P-value (based on LR test): 0.72509

**DE (size-corrected)**

<u>All species individually</u>

Phylogenetic signal lambda: 0.880638
logL(lambda): 44.4041
LR(lambda=0): 27.1398
P-value (based on LR test): **1.89261e-07**

<u>TSG aggregated</u>

Phylogenetic signal lambda: 0.84782
logL(lambda): 20.9816
LR(lambda=0): 5.21661
P-value (based on LR test): **<u>0.0223722</u>**

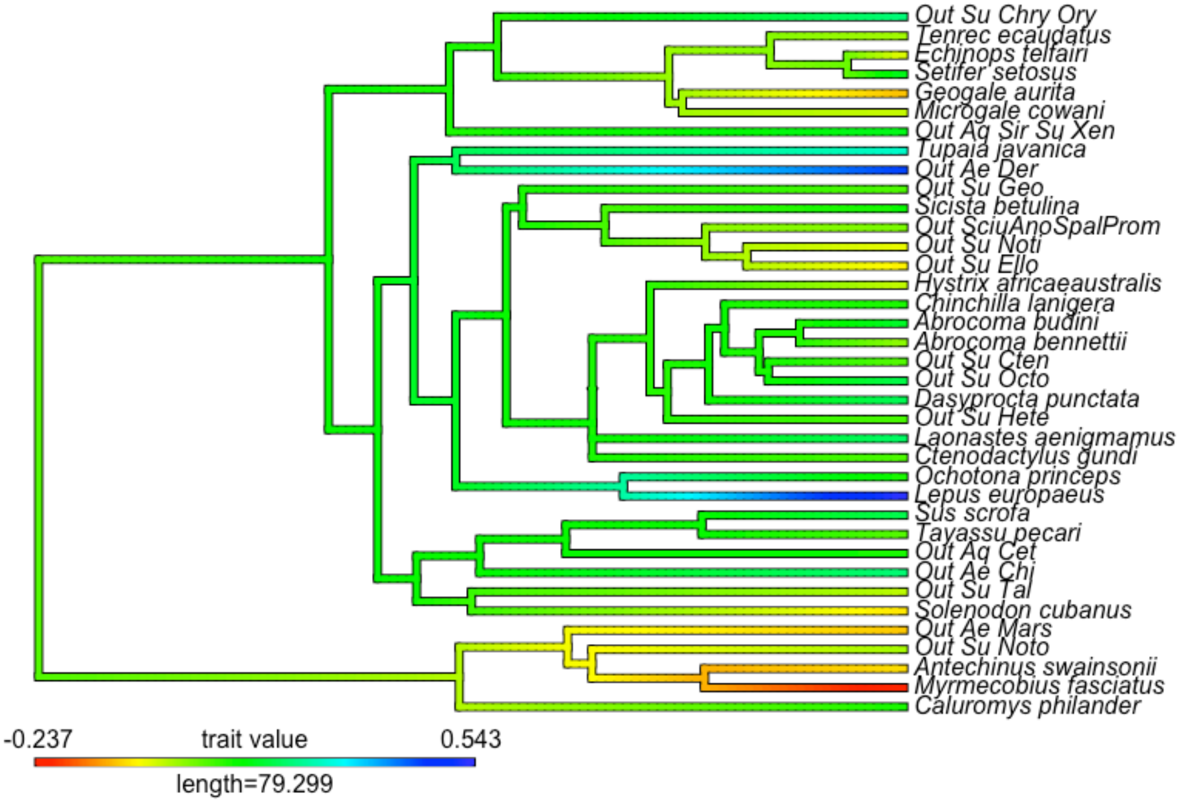

### References


Everson, K. M., K. B. P. Hildebrandt, S. M. Goodman, and L. E. Olson. 2018. Caught in the act: Incipient speciation across a latitudinal gradient in a semifossorial mammal from Madagascar, the mole tenrec Oryzorictes hova (Tenrecidae). Molecular Phylogenetics and Evolution 126:74–84.

Fabre, P., M. Tilak, C. Denys, P. Gaubert, V. Nicolas, E. J. P. Douzery, and L. Marivaux. 2018. Flightless scaly-tailed squirrels never learned how to fly : A reappraisal of Anomaluridae phylogeny. Zoologica Scripta, doi: 10.1111/zsc.12286.

Hothorn, T., F. Bretz, and P. Westfall. 2008. Simultaneous inference in general parametric models. Biometrical Journal 50:346–363.

Kumar, S., G. Stecher, M. Suleski, and S. B. Hedges. 2017. TimeTree: A resource for timelines, timetrees, and divergence times. Molecular biology and evolution 34:1812–1819.

Nowak, R. M. 2020. Walker's Mammals of the World: Monotremes, Marsupials, Afrotherians, Xenarthrans, and Sundatherians. Johns Hopkins University Press, Baltimore.

Revell, L. J. 2012. phytools: An R package for phylogenetic comparative biology (and other things). Methods in Ecology and Evolution 3:217–223.

Springer, M. S., A. V. Signore, J. L. A. Paijmans, J. Vélez-Juarbe, D. P. Domning, C. E. Bauer, K. He, L. Crerar, P. F. Campos, W. J. Murphy, R. W. Meredith, J. Gatesy, E. Willerslev, R. D. E. MacPhee, M. Hofreiter, and K. L. Campbell. 2015. Interordinal gene capture, the phylogenetic position of Steller's sea cow based on molecular and morphological data, and the macroevolutionary history of Sirenia. Molecular Phylogenetics and Evolution 91:178–193.

Wilson, D. E., and D. M. Reeder. 2005. Mammal species of the world: a taxonomic and geographic reference. 3rd ed. John Hopkins University Press, Baltimore.
